# The deubiquitinase OTULIN regulates tau expression and RNA metabolism in neurons

**DOI:** 10.1101/2025.04.09.648063

**Authors:** Karthikeyan Tangavelou, Virginie Bondu, Mingqi Li, Wei Li, Francesca-Fang Liao, Kiran Bhaskar

## Abstract

The degradation of aggregation-prone tau is regulated by the ubiquitin-proteasome system (UPS) and autophagy, which are impaired in Alzheimer’s disease (AD) and related tauopathies causing tau aggregation. Protein ubiquitination with linkage specificity determines the fate of proteins that can be either degradative or stabilization signals. While the linear M1-linked ubiquitination on protein aggregates is a signaling hub that recruits various ubiquitin-binding proteins for coordinated actions of protein aggregates turnover and inflammatory NF-kB activation, the deubiquitinase OTULIN counteracts with the M1-linked ubiquitin signaling. However, the exact role of OTULIN on tau aggregate clearance in AD is unknown. Based on our bulk RNA sequence analysis, human inducible pluripotent stem cell (iPSC)-derived neurons (iPSNs) from an individual with late-onset sporadic AD (sAD2.1) show downregulation of ubiquitin ligase activating factors (MAGEA2B and MAGEA) and OTULIN long non-coding RNA (lncRNA-OTULIN) compared to healthy control WTC11 iPSNs. In sAD2.1 iPSNs, downregulated lncRNA-OTULIN is inversely correlated with increased levels of OTULIN protein and phosphorylated tau at p-S202/p-T205 (AT8), p-T231 (AT180), and p-S396/p-S404 (PHF-1). Loss of OTULIN deubiquitinase function using pharmacological inhibitor UC495 or CRISPR-Cas9-mediated OTULIN gene knockout causes a significant reduction of total tau and phosphorylated tau at AT8 epitope in sAD2.1 iPSNs. Whereas in SH-SY5Y neuroblastoma cells, either treating with the UC495 compound or knocking out of the OTULIN gene causes a significant reduction of total tau at both mRNA and protein levels and consequently decreases phosphorylated tau at AT8, AT180, and PHF-1 epitopes. An additional bulk RNA sequence analysis of OUTLIN knockout SH-SY5Y shows a 14-fold down-regulation of tau mRNA levels and differential expression of many other genes associated with autophagy, UPS, NF-kB pathway, and RNA metabolism. Together, our results suggest for the first time a non-canonical function for OTULIN in regulating gene expression and RNA metabolism, which may have a significant pathogenic role in AD and related tauopathies.

## Introduction

The tau is encoded by the microtubule-associated protein tau or *MAPT* gene, which is located on chromosome 17q21 and expressed as six isoforms by alternative splicing in the human brain^1,2^. Tauopathy is a group of more than 20 neurodegenerative diseases, including Alzheimer’s disease (AD), caused by intracellular aggregation of hyperphosphorylated tau as neurofibrillary tangles (NFTs) in neurons^3,4,5^. Lack of improper clearance of misfolded/hyperphosphorylated tau/NFTs leads to its aggregation in AD and other tauopathies. Depending on the context in the neurons, tau can be a substrate of 20S proteasome, 26S proteasome, chaperone-mediated autophagy, microautophagy, macroautophagy, and aggrephagy (reviewed elsewhere^6,7^). Prior studies have demonstrated that impairment in these tau-clearing mechanisms can lead to tau accumulation as NFTs in tauopathy brains^7^.

The linear ubiquitin assembly complex (LUBAC) (Ring finger protein 31/HOIL-1-interacting protein (RNF31/HOIP), RanBP-type and C3HC4-type zinc finger-containing protein 1/Heme-oxidized IRP2 ubiquitin ligase 1 (RBCK1/HOIL-1), SHANK associated RH domain interactor (SHARPIN)) is the only known E3 ubiquitin ligase responsible for adding M1-linked linear ubiquitin chains to protein aggregates^8^. M1-linked ubiquitin is enriched around the nucleus and spread throughout the cell body of a neuron in various proteinopathies, including AD, frontotemporal dementia (FTD), Parkinson’s disease (PD), polyglutamine (polyQ) diseases, and amyotrophic lateral sclerosis (ALS)^10, 9^. The ubiquitin chaperone, valosin-containing protein (VCP/p97 ATPase) recruits RNF31/HOIP to the sites of polyQ aggregates in Huntington’s disease (HD) models for efficient clearance of Huntingtin (Htt)-polyQ aggregates by recruiting autophagy receptors via M1-linked ubiquitin chains^10^. In another study, lysine (K)48-linked ubiquitination on tau aggregates is a prerequisite for recruiting VCP, which interacts with HSP70 to extract tau from aggregates for proteasomal degradation or extracellular secretion of seeding competent tau^10^. Moreover, the tandem mass spectrometry analysis of tau-paired helical filaments (PHFs) isolated from the autopsy of human AD brains shows the presence of various ubiquitin chains, including methionine (M)1-, K6-, K11-, K48-, and K63-linked ubiquitination^11,12,13^.

M1-linked ubiquitination on protein aggregates and host-invaded microbes is a signal for activation of nuclear factor-kappa B cells (NF-κB), which regulates the transcription of DNA associated with immune response and cell survival under various stresses and infection^14,15,16,17,18,19,20,21^. Most often, neuronally derived protein aggregates such as NFTs, α-synuclein, and Htt-polyQ are inflammatory upon the addition of M1-linked ubiquitin chains, provide a binding platform to NF-κB regulator/autophagy adaptor, NF-κB essential modifier (NEMO), and autophagy receptors for NF-κB activation and protein aggregates clearance, respectively^10,11,22,23^. Various other ubiquitin-specific linkages in NFTs recruit proteasome and aggrephagy machinery for tau clearance or extracellular secretion of seeding competent tau upon impairment of proteostasis^8,12^.

The ovarian tumor (OTU) domain-containing deubiquitinase with linear linkage specificity (OTULIN) is exclusively the interlinear M1-linked ubiquitin chains but not the peptide bond between target proteins and the first ubiquitin ^24^, whereas cylindromatosis (CYLD) deubiquitinates both M1- and K63-linked ubiquitin chains^25,26,27^. Both deubiquitinases OTULIN and CYLD bind to HOIP/RNF31 and decrease LUBAC-mediated M1-linked ubiquitination on target proteins and consequently decreases NF-κB activation under various inflammatory stimuli^28,29,30,31,32^. Despite their role in negative regulation of NF-κB activity, OTULIN knock-out show increased levels of M1-linked ubiquitination on LUBAC components, RBCK1/HOIL-1 and SHARPIN, but not with CYLD knockout suggesting that OTULIN directly regulates LUBAC activity in basal condition^29^. Moreover, OTULIN tyrosine (Y) 56 is essential for bonding with LUBAC component, HOIP/RNF31 prevents TNFα-induced NF-κB activation, whereas phospho-OTULIN Y56 prevents interaction with HOIP and activates NF-κB^31,33^ In another study, OTULIN deubiquitinates LUBAC itself polyubiquitinated M1-linked linear chains and prevents LUBAC-mediated cell death and inflammation^34^.

Mice knockout for *HOIP*^35^, *HOIL-1*^36^, and *OTULIN*^35^ genes are embryonic lethal, whereas SHARPIN deficiency causes chronic autoinflammatory disease in the skin of mice ^37^. Mutations in human HOIP L72P/Q399H^38,39^ and HOIL-1 deficiency^40^ cause immunodeficiency and autoinflammation due to a lack of LUBAC-mediated M1-linked ubiquitination. Whereas mutations in human OTULIN Y244C/L272P/G281R decrease deubiquitinase activity, resulting in increased levels of M1-linked polyubiquitination in OTULIN-related autoinflammatory syndrome (ORAS) patients^41,42,43^. The effect of OTULIN deficiency varies among cell types, such as myeloid macrophages that cause elevated autoinflammation associated with increased M1-linked polyubiquitination and NF-κB activity. *OTULIN* deficiency in lymphoid B or T cells causes no adverse inflammation due to complete loss of LUBAC components, HOIP, and SHARPIN at protein levels by proteasomal activity, but not their mRNAs^44^. Despite the deubiquitinase, OTULIN negatively regulates inflammation and autoimmunity by counteracting with LUBAC-mediated linear M1-linked polyubiquitination, which is essential to recruit autophagy adaptors and receptors for efficient clearance of inflammatory stimuli. OTULIN also functions as negative and positive regulators of autophagy^44^ and proteasomes^45^, respectively.

Here, we describe the role of OTULIN in dysregulating pathological tau aggregates clearance in human induced pluripotent stem cells (iPSCs)-derived neurons (iPSNs) from sporadic AD, sAD2.1 line^46^. We identified that the expression of many genes downregulated, including the ubiquitin ligase activating factor melanoma-associated antigen (MAGE) family members (MAGE-A2B and MAGE-A), and upregulation of long non-coding RNA (lncRNA) for OTULIN in sAD2.1 neurons. Increased OTULIN level is positively correlating with tau pathology markers such as increased phosphorylated tau at p-S202/p-T205 (AT8), p-T231 (AT180), and p-S396/p-S404 (PHF-1) in sAD2.1 neurons. Interfering OTULIN function by inhibiting its deubiquitinase activity with UC495 compound or CRISPR-Cas9-mediated OTULIN deficiency in sAD2.1 neurons decreases total tau and phosphorylated tau levels. Surprisingly, OUTLIN knockout in human neuroblastoma SH-SY5Y line shows complete loss of mRNA transcripts of *MAPT*, *HOIP*, *HOIL-1*, *SHARPIN*, and regulators of NF-κB and autophagy. Our initial thought of OTULIN deficiency leading to clearance of pathological tau (p-Tau) is overshadowed by the master-regulatory role of OTULIN in RNA metabolism. This non-canonical novel function of OTULIN in neurons suggests its potential role in RNA metabolism in neurodegenerative diseases.

## Results

### OTULIN and phosphorylated tau levels are elevated in sporadic sAD2.1 iPSC line-derived neurons

We performed bulk RNA sequence analyses in sporadic Alzheimer’s Disease (sAD2.1) induced pluripotent stem cell line (iPSC)-derived neurons (iPSNs) and healthy control WTC11 iPSNs to identify the differentially expressed genes. Details about the mapped region, sequence coverage, ready density, principal component analysis, Pearson correlation matrix, and significantly altered biological processes/pathways are shown in Supplementary Figures S1-S8 and Table S1-S2. Our RNA sequencing data shows that 2,390 and 2,124 genes were significantly up- or down-regulated in sAD2.1 iPSNs as compared to healthy control WTC11 neurons, respectively (Supplementary Fig. S5). Similarly, there were 3,828 significantly upregulated and 1,852 down-regulated transcripts were noted in sAD2.1 neurons as compared to WTC11 neurons (Supplementary Fig. S5). Gene Ontology (GO) mapped >2,000 significantly altered genes to be associated with ‘protein binding’ as a major molecular functional pathway (Supplementary Fig. S6). The GO and Kyoto Encyclopedia of Genes and Genomes (KEGG) enrichment analyses show that the regulators of ubiquitin ligase activity such as MAGE family members (*MAGEA2B* and *MAGEA*) – that are known to increase ubiquitin ligase activity were down-regulated in sAD2.1 neurons as compared to WTC11 neurons (Fig. 1A-B). As expected, there was a positive correlation between sAD2.1 with AD and tauopathies in Gene set enrichment analyses (GSEA) (Fig. 1C-D). Conversely, the WTC11 was negatively correlated with these two conditions. A notable alteration was observed in long non-coding RNA (lncRNA) derived from *OTULIN* gene. LncRNA-OTULIN was significantly downregulated in sAD2.1 neurons compared to WTC11 neurons (Fig. 1E). Conversely, the OTULIN protein level was significantly elevated in sAD2.1 than in WTC11 neurons (Fig. 1F-G). In addition to an increased level of OTULIN, tau phosphorylated at p-S202/p-T205 (AT8), p-T231 (AT180), and p-S396/p-S404 (PHF-1) was also significantly elevated in sAD2.1 neurons (Fig. 1F-G).

**Figure 1:**
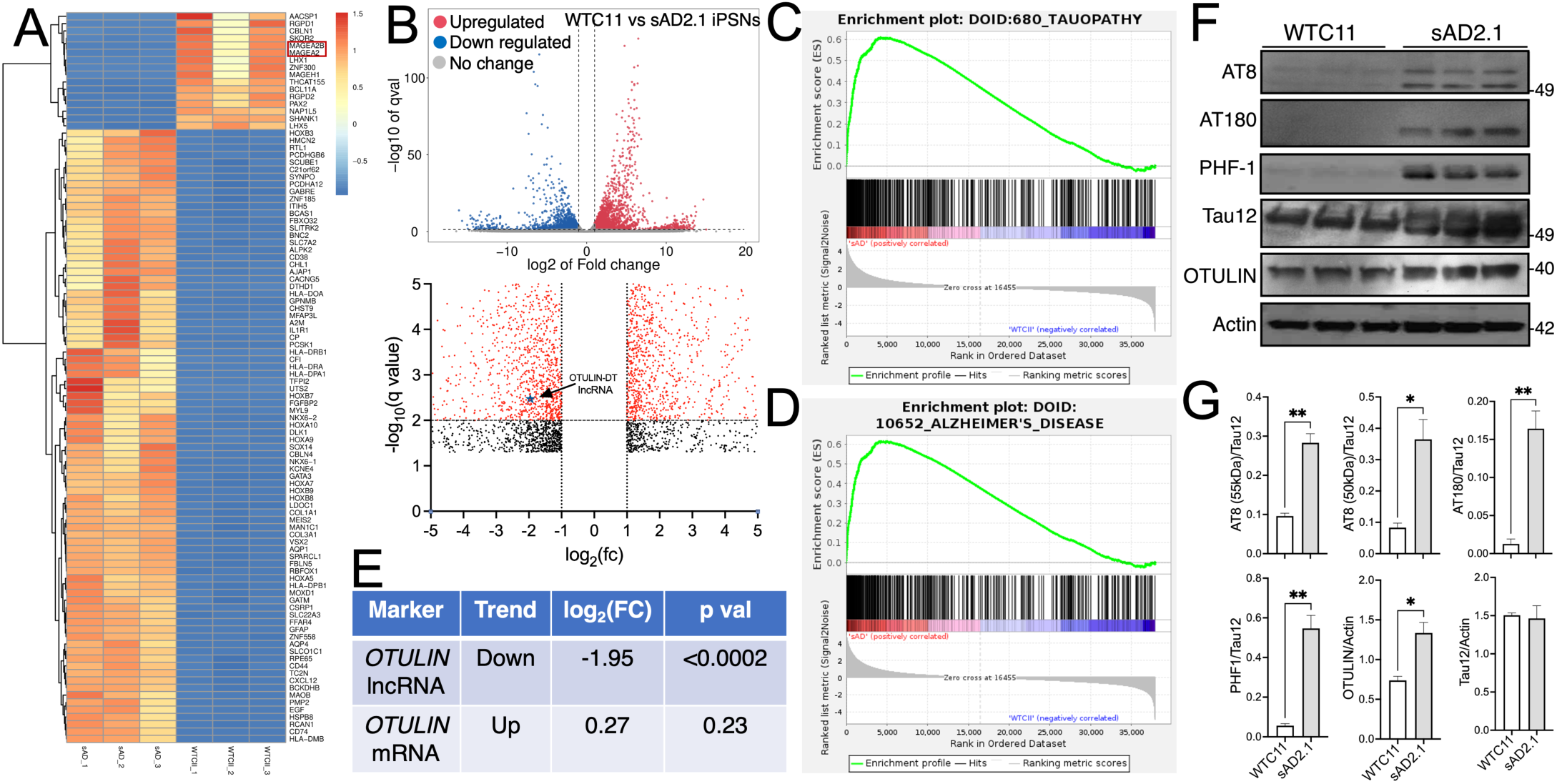
Sporadic sAD2.1 iPSC line-derived neurons show significantly altered gene expression profile, elevated OTULIN, and hyperphosphorylated tau levels compared to healthy control WTC11 neurons. (A) Heat-map showing differentially expressed mRNA levels in sAD2.1 versus WTC11 neurons. Note that the MAGE Family Member A2B (*MAGEA2B* and *MAGEA*), which are known to enhance ubiquitin ligase activity, is significantly reduced in sAD2.1 compared to WTC11 neurons. (B) Volcano plots showing the numbers of up- and down-regulated genes in sAD2.1 neurons as compared to WTC11 control. (C) Gene Set Enrichment Analyses (GSEA) pathways showing enrichment of signaling pathways relevant to tauopathy and Alzheimer’s disease. (E) Otulin lncRNA was one of the significantly altered transcripts (downregulated with log2 fold change of −1.95; p<0.0002) and conversely, slight upregulation of OTULIN mRNA. (F-G) Western blot analyses showing significantly elevated phosphorylated tau at p-S202/p-T205 (AT8), p-T231 (AT180), and p-S396/p-S404 (PHF-1) and OTULIN in sAD2.1 as compared to healthy control WTC11 neurons. Data presented as mean ± SEM; unpaired *t* test; *p<0.05; **p<0.01; n=3.

### Pharmacological inhibition of OTULIN deubiquitinase with compound UC495 significantly decreases phosphorylated tau p-S202/p-T205 (AT8) levels in sporadic sAD2.1 iPSC line-derived neurons

The deubiquitinase OTULIN level is increased in sporadic sAD2.1 iPSC-derived neurons (Fig. 1F-G) that can potentially increase its deubiquitinase activity towards the linear M1-linked ubiquitin chains and consequently increase tau pathology. To test this hypothesis, sAD2.1 neurons were treated with compound UC495 to inhibit OTULIN deubiquitinase activity (Supplementary Fig. S14) and assessed tau pathology. Inhibiting OTULIN deubiquitinase activity significantly reduced the level of phosphorylated tau at p-S202/p-T205 (AT8), and modestly decreased p-S396/p-S404 (PHF-1) positive tau levels compared to vehicle-treated sAD2.1 neurons. However, inhibiting the deubiquitinase activity of OTULIN with UC495 compound did not affect the levels of either OTULIN or total tau (Fig. 2A-B)

**Figure 2:**
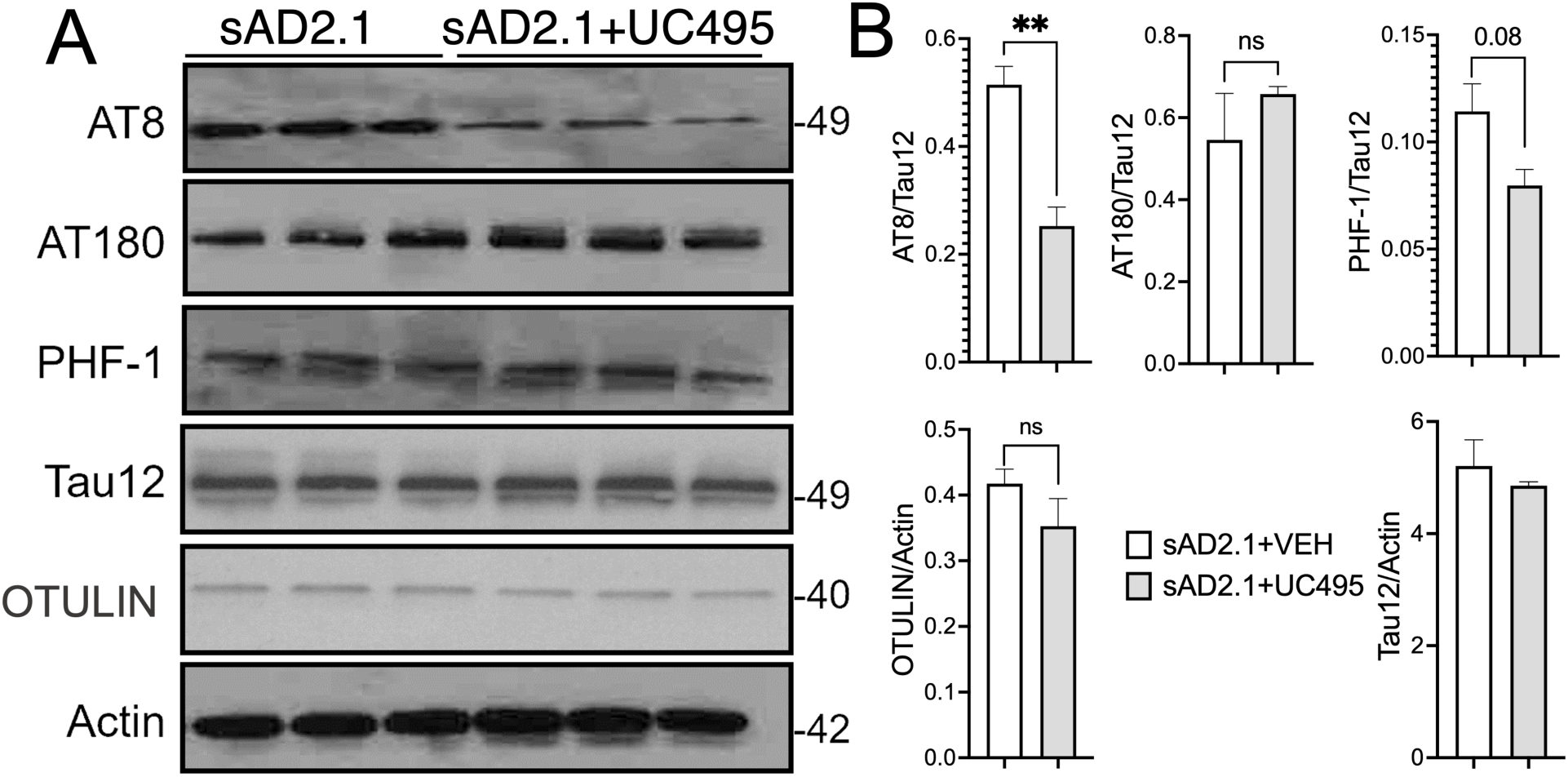
Pharmacological inhibition of OTULIN deubiquitinase with compound UC495 selectively decreases phosphorylated tau at p-S202/p-T205 (AT8) in sAD2.1 iPSNs. (A-B) Western blot analyses showing significantly decreased phosphorylated tau at p-S202/p-T205 (AT8), and a modest reduction of tau p-S396/p-S404 (PHF-1) upon pharmacological inhibition of OTULIN with compound UC495 in sAD2.1 iPSC line-derived neurons. Data presented as mean ± SEM; unpaired *t* test; **p<0.01; n=3.

### CRISPR-Cas9-mediated knockout of OTULIN gene significantly decreases total tau in sAD2.1 iPSC line-derived neurons

Unlike pharmacological inhibition of OTULIN deubiquitinase activity with compound UC495, CRISPR-Cas9-mediated knockout of *OTULIN* gene completely abolished its expression in sAD2.1 iPSC line-derived neurons (Fig. 3A). As a result, both total and phosphorylated tau levels significantly decreased in OTULIN deficient sAD2.1 neurons (Fig. 3A-B). Since the complete loss of tau in OTULIN deficient sAD2.1 neurons, we wondered whether differentiated neurons retained the neuronal marker, NeuN, which confirms no adverse effect on neuronal differentiation despite the absence of both OTULIN and tau (Fig. 3A-B). We also imaged the differentiated neurons using the bright field microscope, which showed no significant morphological difference or alterations in NeuN levels noted either in the presence or absence of *OTULIN* sAD2.1 iPSC-derived neurons (Fig. 3C). Together, these results suggest that CRISPR-Cas9-mediated *OTULIN* knockout in sporadic sAD2.1 iPSC line-derived neurons causes complete loss of total tau and phosphorylated tau levels without affecting neuronal survivability.

**Figure 3:**
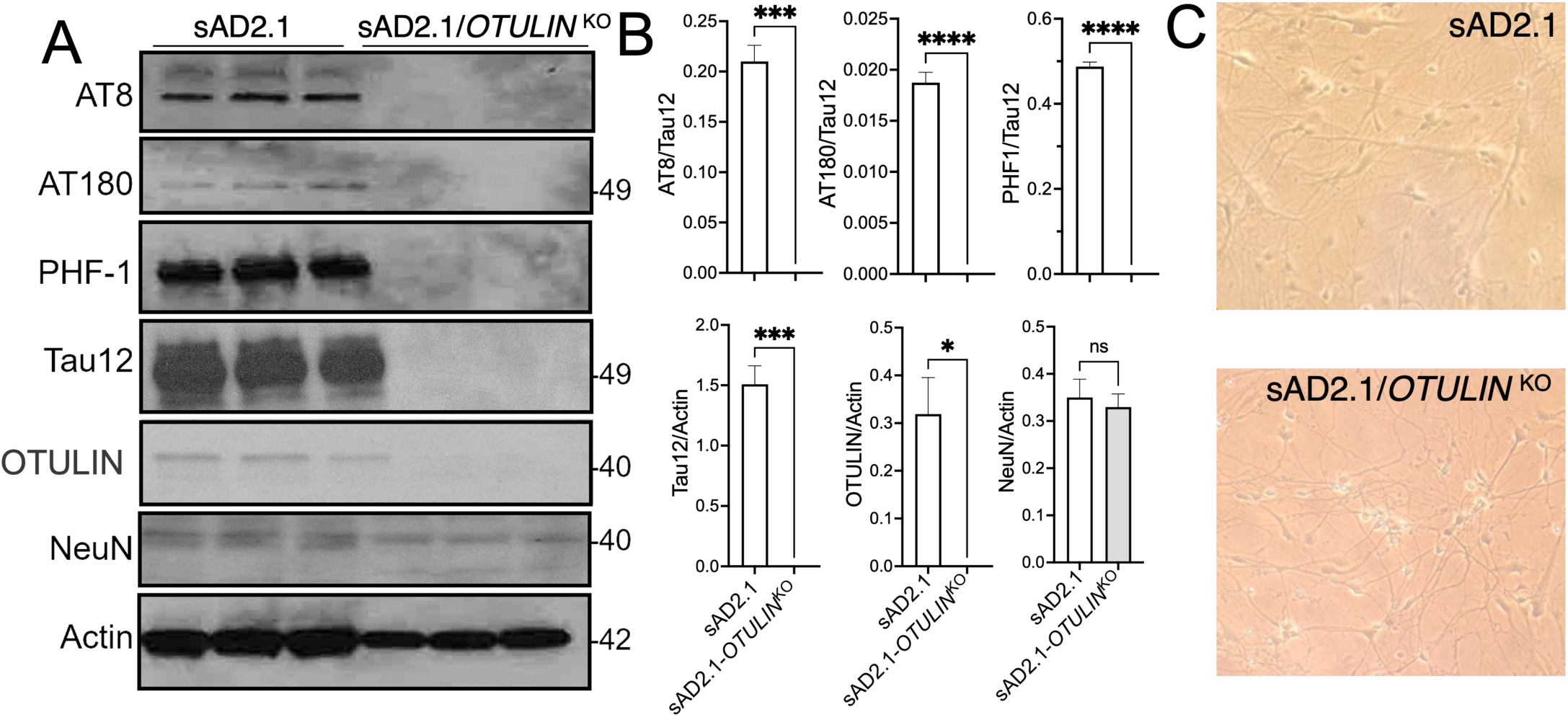
CRISPR-Cas9-mediated knockout of OTULIN causes a complete loss of total tau in sporadic sAD2.1 neurons. (A-B) Western blot quantifications showing significantly reduced phosphorylated tau at p-S202/p-T205 (AT8), p-T231 (AT180), and p-S396/p-S404 (PHF-1) as well as total tau in CRISPR-Cas9-mediated OTULIN KO of sporadic sAD2.1 iPSC line-derived neurons as compared to sAD2.1 neurons. Note that the reduction is not due to toxicity to neurons, as no significant difference in the NeuN levels were noted. (C) Representative images from light microscopy showing the live matured sAD2.1 iPSC line-derived neurons as control vs. sAD2.1 OTULIN KO iPSC line-derived neurons. Data presented as mean ± SEM; unpaired *t* test; *p<0.05; **p<0.01; ***p<0.005; ****p<0.001; n=3.

### Inhibiting OTULIN deubiquitinase with compound UC495 significantly decreases the levels of OTULIN, total tau, and ubiquitin in the SH-SY5Y cell line

In contrast to the effect of UC495 in modestly reducing p-S202/p-T205 (AT8) tau in sporadic sAD2.1 iPSNs, inhibiting deubiquitinase activity of OTULIN with compound UC495 in SH-SY5Y significantly decreased the levels of OTULIN, ubiquitin, and total tau and consequently decreases phosphorylated tau at p-S202/p-T205 (AT8), p-T231 (AT180), p-S396/p-S404 (PHF-1) (Fig. 4A-B). Despite reducing endogenous tau levels, we did not observe cell toxicity as the actin level was relatively uniform between UC495-treated and control SH-SY5Y cells (data not shown).

**Figure 4:**
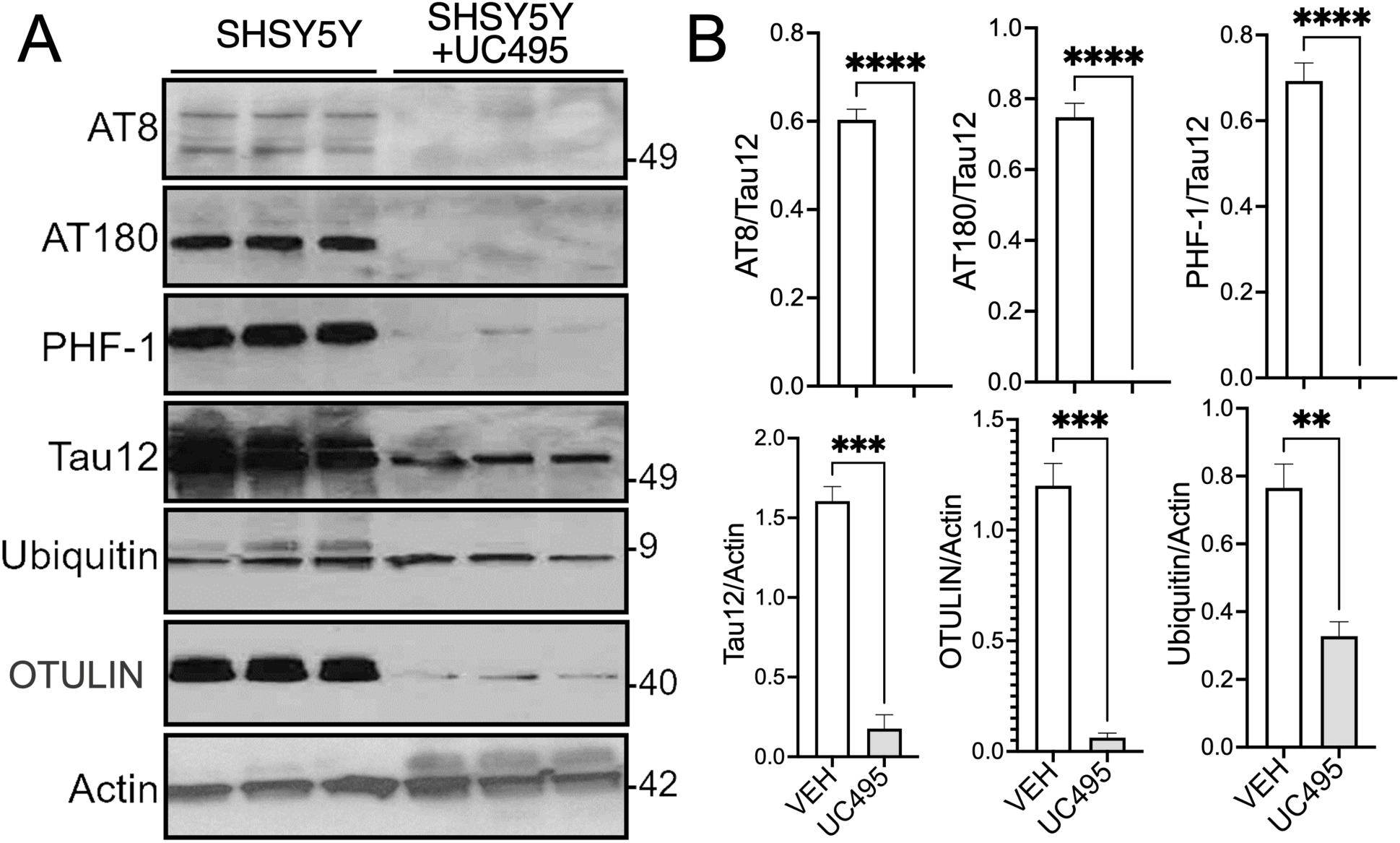
Pharmacological inhibition of OTULIN with UC495 compound significantly decreases hyper-phosphorylated and total tau, OTULIN and ubiquitin in SH-SY5Y cells. (A-B) Western blot quantifications showing significantly reduced phosphorylated tau at p-S202/p-T205 (AT8), p-T231 (AT180), and p-S396/p-S404 (PHF-1) as well as total tau upon inhibition of OTULIN deubiquitinase with UC495 compound in SH-SY5Y human neuroblastoma line as compared to DMSO treated control. OTULIN inhibition also decreases OTULIN and ubiquitin levels. Data presented as mean ± SEM; unpaired *t* test; **p<0.01; ***p<0.005; ****p<0.001; n=3.

### CRISPR-Cas9-mediated OTULIN knockout significantly decreases total tau and ubiquitin in SH-SY5Y

Similar to the pharmacological inhibition of OTULIN deubiquitinase with compound UC495, the CRISPR-Cas9-mediated gene interfering with *OTULIN* completely abolished its protein expression in the SH-SY5Y line (Fig. 5A). Absence of OTULIN induces a significant reduction of total tau. Consequently, it decreases the level of phosphorylated tau (Fig. 5A-B). Moreover, the effect of OTULIN deficiency is also reflected in the level of total ubiquitin, whose level is significantly decreased, similar to inhibiting the deubiquitinase activity of OTULIN with UC495 compound (Fig. 5A-B). Collectively, inhibiting deubiquitinase of OTULIN with small molecule UC495 or CRISPR-Cas9-mediated knockout of *OTULIN* gene in the SH-SY5Y line decreases OTULIN levels, which in turn significantly reduces the levels of total tau and ubiquitin.

**Figure 5:**
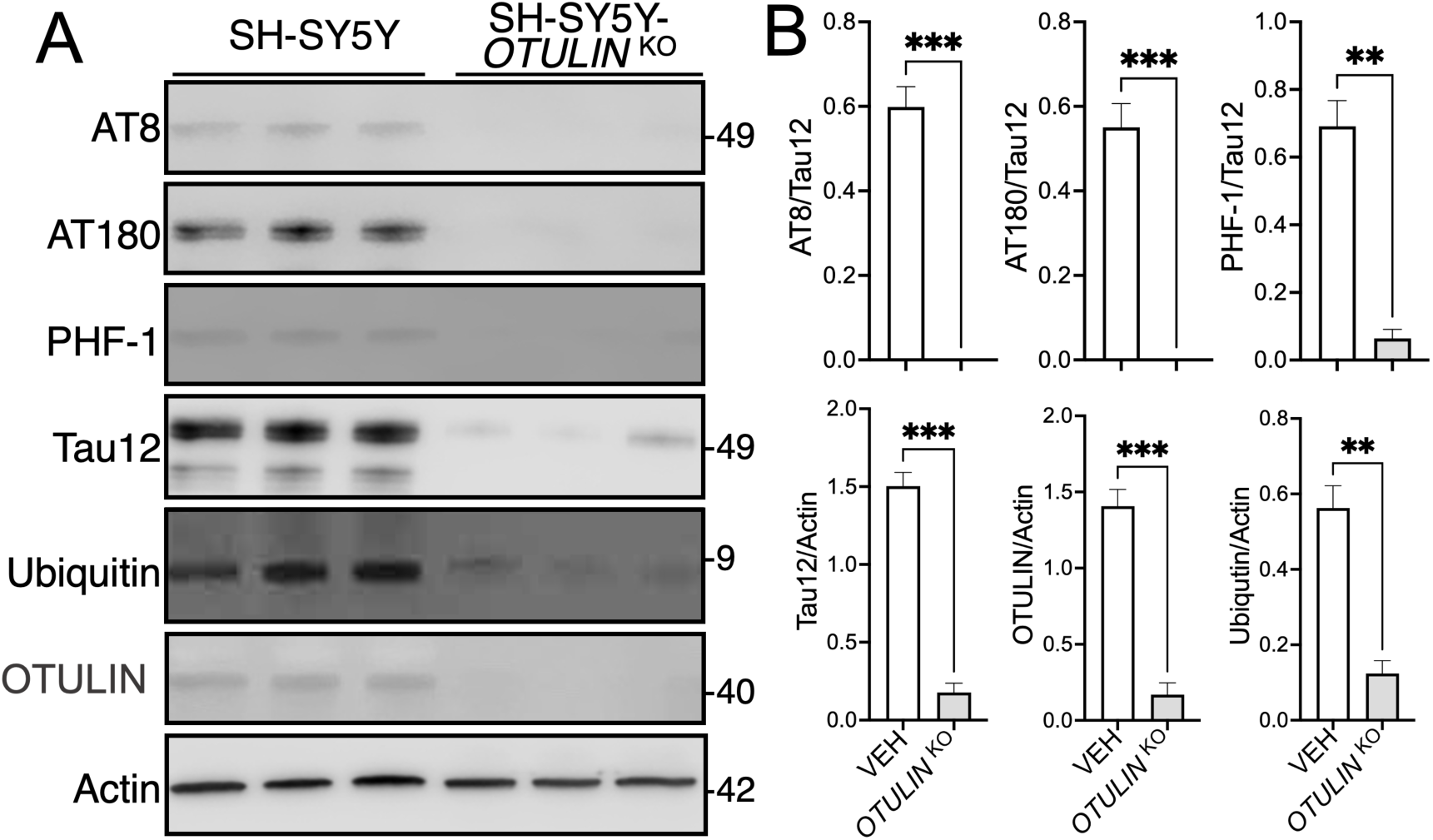
CRISPR-Cas9-mediated *OTULIN* knockout significantly decreases total and hyperphosphorylated tau and ubiquitin in SH-SY5Y human neuroblastoma cells. (A-B) Western blot quantifications showing significantly reduced phosphorylated tau at p-S202/p-T205 (AT8), p-T231 (AT180), and p-S396/p-S404 (PHF-1) as well as total tau in CRISPR-Cas9-mediated *OTULIN* KO SH-SY5Y cells as compared to control SH-SY5Y wild type (WT). *OTULIN* KO also significantly decreases ubiquitin levels. Data presented as mean ± SEM; unpaired *t* test; **p<0.01; ***p<0.005; ****p<0.001; n=3.

### None of the major proteostasis inhibitors restores tau levels in OTULIN-deficient SH-SY5Y line

We predicted that the absence of OTULIN deubiquitinase in the SH-SY5Y line might enhance tau degradation by proteostasis activity. To determine proteostasis activity causing degradation of tau in SH-SY5Y *OTULIN* KO, cells were pretreated with protein translation inhibitor, cycloheximide (CHX) for 1h and added various proteostasis inhibitors individually or together for additional 4h with lactacystin (20S proteasome), MG132 (26S proteasome), Bafilomycin A1 (Baf A1; autophagy) and PD 150606 (Calpain 1/2). None of the inhibitors restored tau level in OTULIN deficient SH-SY5Y cells (Fig. 6A), which motivated us to investigate the presence of tau at *MAPT* mRNA level by Real-time quantitative reverse transcription PCR (Real-time-RT-qPCR) along with 18S ribosomal RNA (18s rRNA) as a housekeeping reference gene for normalizing RT-qPCR data. Surprisingly, *MAPT* mRNA was undetectable in the OTULIN-deficient SH-SY5Y line compared to the wild-type line (Fig. 6B).

**Figure 6:**
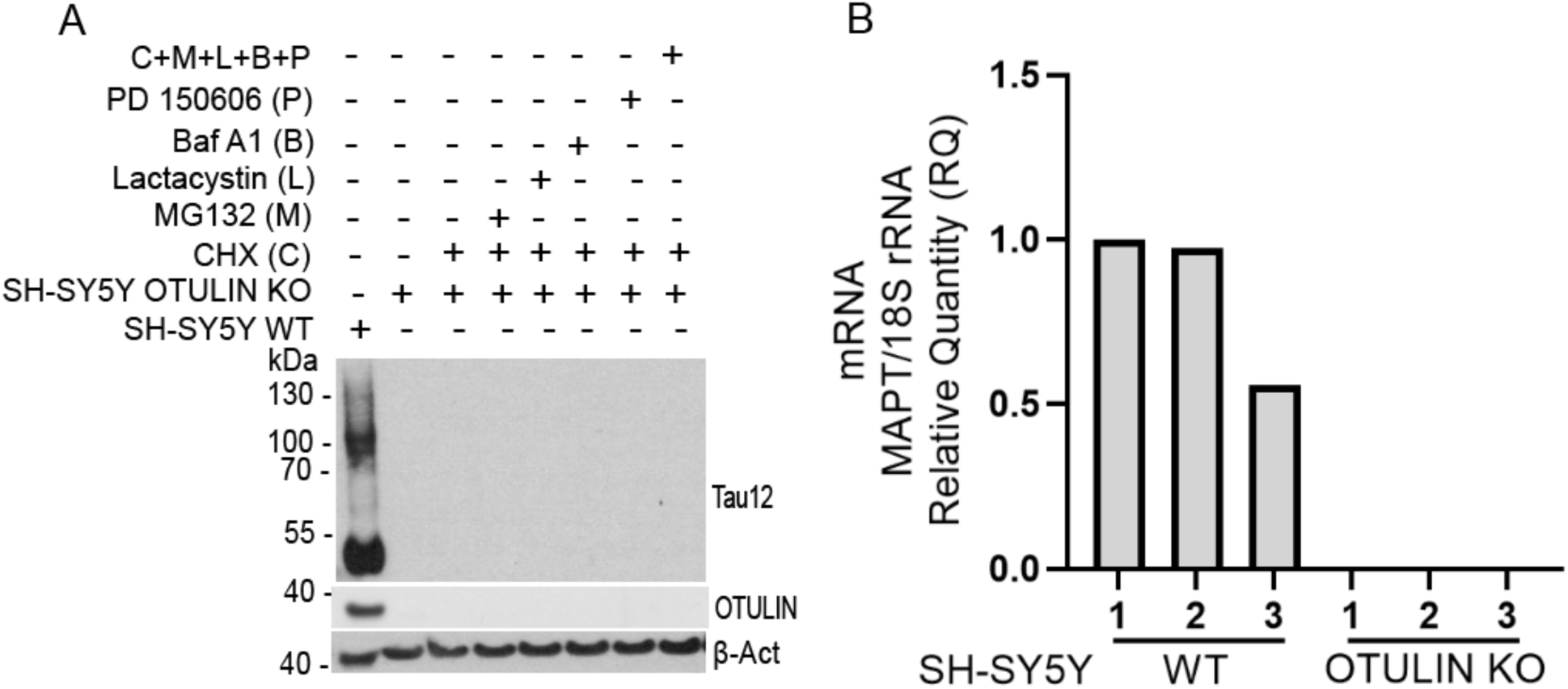
OTULIN deficiency in SH-SY5Y downregulates tau encoding *MAPT mRNA*. (A) Protein translation was inhibited with 50µM cycloheximide (CHX) 1h prior adding the proteostasis inhibitors, 10µM MG132 (26S proteasome), 10µM lactacystin (20S proteasome), 100nM bafilomycin A1 (Baf A1, autophagy), 100µM PD 150606 (calpain 1/2) either individually or all four inhibitors together for 4h. None of the inhibitors stabilizes tau in the absence of OTULIN. (B) Real-time quantitative reverse transcription PCR (Real-time-RT-qPCR) analysis shows the absence of *MAPT* mRNA in *OTULIN* knockout (KO1/2/3) compared to wild type (WT1/2/3) SH-SY5Y. 18S ribosomal RNA (18s rRNA) was used as a housekeeping reference gene for normalizing RT-qPCR data.

### OTULIN regulates RNA metabolism in human SH-SY5Y line

To determine whether OTULIN deficiency has any global effects on the mRNAs of other genes or if it is specific to *MAPT* mRNA, we performed bulk RNA sequencing analyses. We isolated total RNA from SH-SY5Y *OTULIN* KO and wild-type (WT) cells. We found that 774 genes were significantly upregulated and 13,341 genes were downregulated in the OTULIN KO compared to the WT line (Fig. 7A-B). To further confirm that OTULIN regulates RNA metabolism, we performed RNA transcript analyses and identified the upregulation of 1,113 transcripts and downregulation of 43,003 transcripts in the *OTULIN-*deficient SH-SH5Y line (Fig. 7C-D). To classify the differentially expressed genes belongs to specific functional characteristics, we performed a gene ontology (GO) enrichment analysis, which showed differentially expressed genes are mainly associated with protein, nucleotide and RNA binding functions, ribonucleoprotein complex, transferase activity, DNA repair and mitochondrial proteins (Fig. 7E). The Kyoto encyclopedia of genes and genomes (KEGG) cellular pathway enrichment analysis revealed that most of the differentially expressed genes belongs to RNA degradation, RNA polymerase, neurodegeneration, lysosome, autophagy, oxidative phosphorylation, nucleotide excision repair and N-Glycan biosynthesis pathways (Fig. 7F). Heatmap comparative analysis showed a few highly significantly up (yellow, orange, red) and downregulated (blue) gene expression from both *OTULIN* KO and WT (Fig. 8A). Notably, the tau encoding *MAPT* gene transcription is downregulated along with OTULIN interaction partners predicted by STRING protein-protein interaction network (Fig. 8B). Since OTULIN is deubiquitinase and dysregulates RNA metabolism, we mainly focused on the expression level of ubiquitin-activating enzyme (E1), ubiquitin-conjugating enzyme (E2) and RNA-binding ubiquitin ligase enzyme (E3), and RNA decay/degradation factors. OTULIN deficiency downregulated ubiquitin B/C and upregulated E1 enzyme UBA3, many E2 conjugating enzymes, and RNA-binding ubiquitin ligases, *RC3H2* and *MEX3C* (Fig. 8C). *OTULIN* KO also upregulated many RNA degradation/decay-associated proteins, heterogeneous ribonucleoproteins, and a few neurodegenerative disease-associated RNA-binding proteins (Fig. 8D). Details about mapped region, sequence coverage, ready density, principal component analysis, Pearson correlation matrix, and significantly altered biological processes/pathways are shown in Supplementary Figures S9-S13 and Table S3-S4.

**Figure 7:**
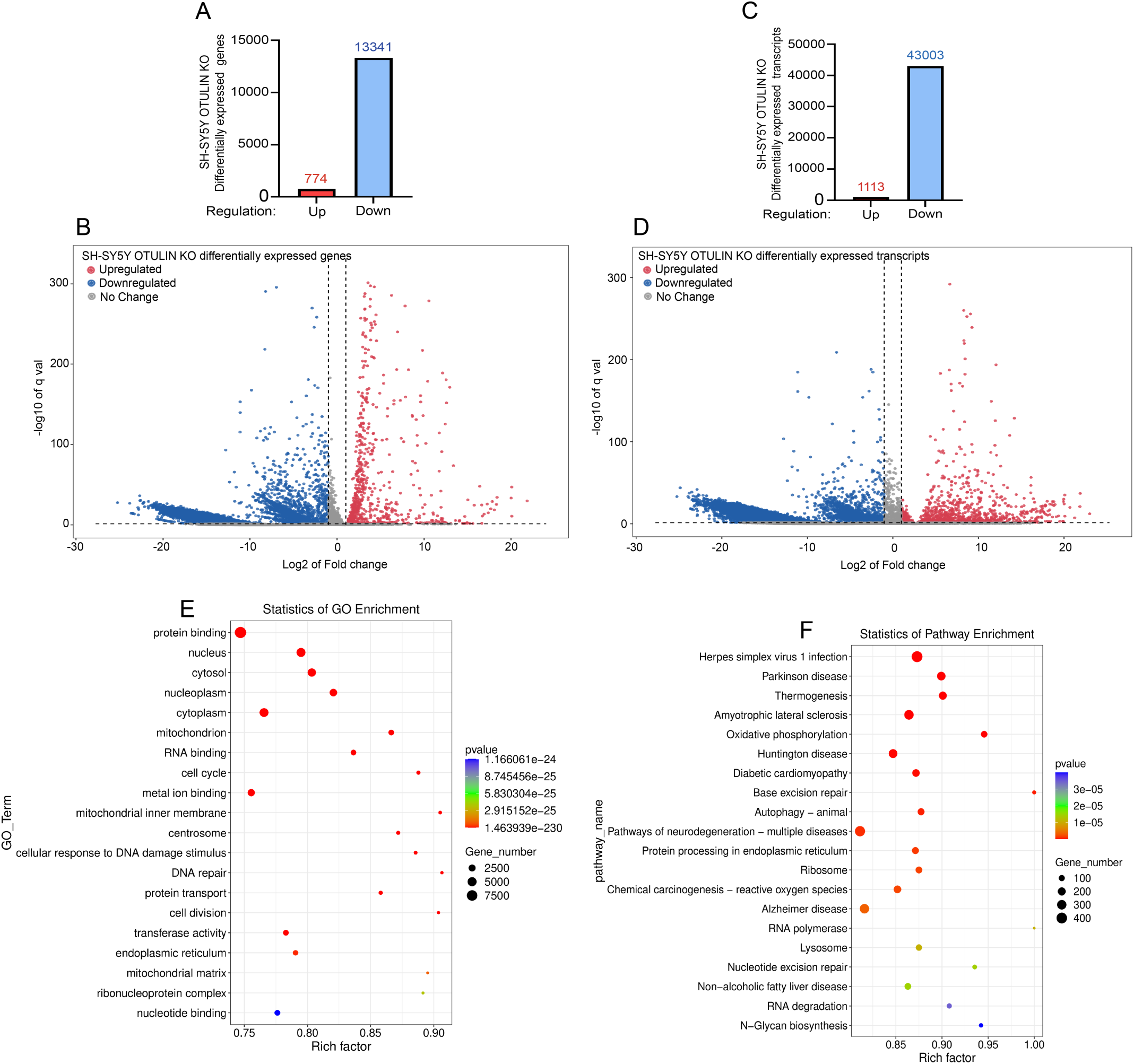
OTULIN is a master regulator of gene expression and RNA metabolism. (A) Bulk RNA sequence analysis of *OTULIN* deficient (KO) SH-SY5Y neuroblastoma shows upregulation of 774 genes and downregulation of 13,341 gene expression compared to wild type (WT) with three replicates from each group. (B) Gene expression volcano plot represents the log2 of the fold change of differentially expressed genes in *OTULIN* KO line with a q value of <0.05 as significant. In log2 of fold changes, red dots at positive and blue dots at negative zones represent up and downregulated gene expression, respectively. Whereas gray dots at the neutral zone denote no significant change of gene expression in both WT and KO lines. (C) OTULIN deficiency dysregulates RNA metabolism as reflected in the number of differentially expressed transcripts, with a downregulation of 43,003 transcripts and an upregulation of 1,113 transcripts as compared to WT. (D) RNA transcript volcano plot represents the log2 of fold change with q value of <0.05 as the significant cutoff for up (red dots) or downregulated (blue dots) and no change (gray) of transcripts level in *OTULIN* KO as compared to WT. (E) A gene ontology (GO) scatter plot represents a set of genes that are associated with protein-to-nucleotide binding functions that were significantly differentially expressed in the *OTULIN* KO line. (F) The cellular pathway analysis using Kyoto Encyclopedia of Genes and Genomes (KEGG) scatter plot describes the enrichment of differentially expressed genes in *OTULIN* KO. RNA metabolism and neurodegenerative disease-associated pathway genes were enriched among the differentially expressed genes. The size and color of dots on the scattered plots (E, F) denote the number of genes and levels of significantly enriched differentially expressed genes.

**Figure 8:**
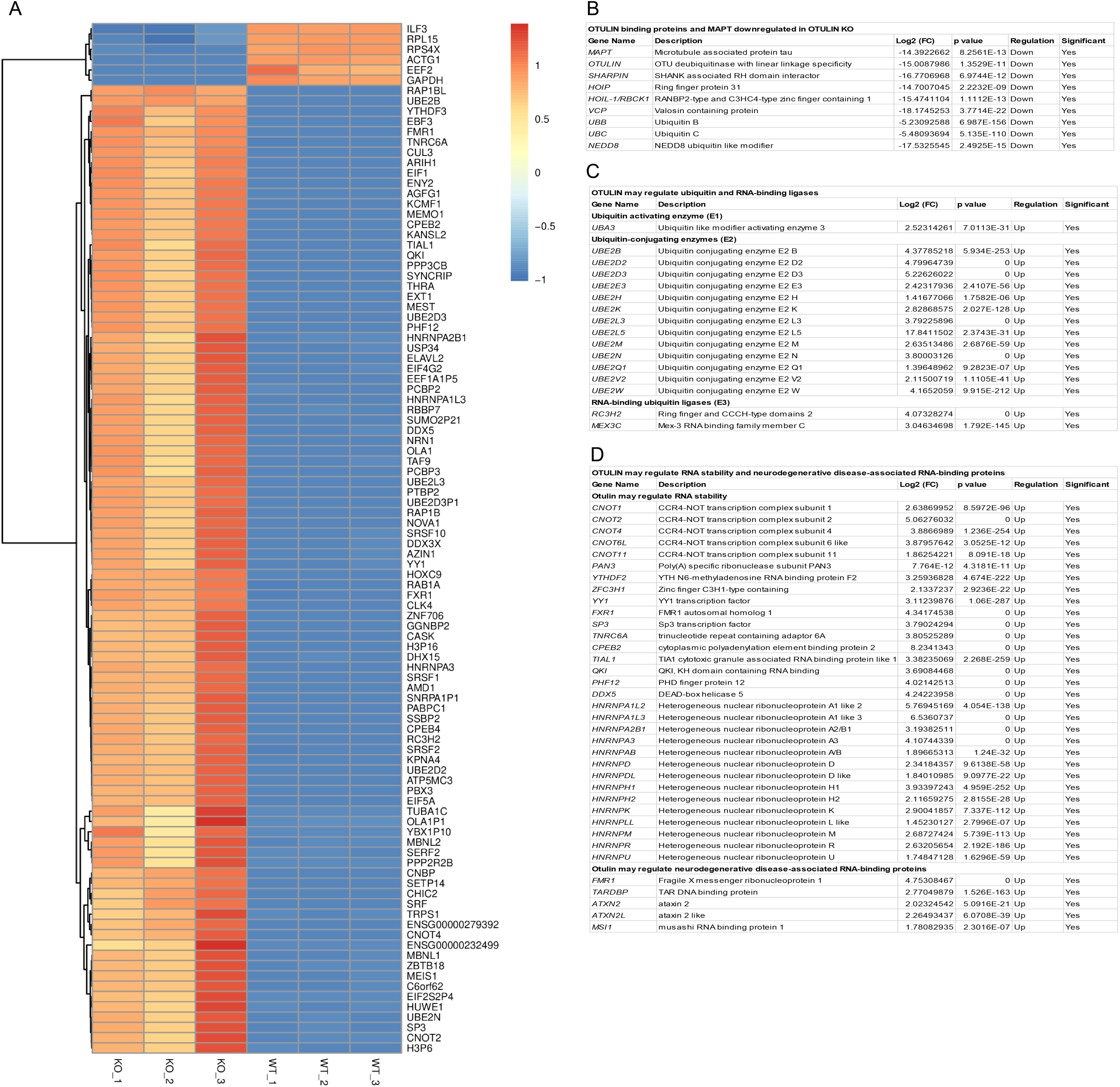
OTULIN deficiency negatively regulates RNA metabolism. (A) Heatmap analysis of differentially expressed genes from *OTULIN* KO (KO_1,2,3) and wild type (WT_1,2,3) SH-SY5Y lines. Heatmap intensity in color codes, orange to red and blue, indicates up and downregulated gene expression, respectively. Most of the upregulated gene expression in *OTULIN* KO is linked with mRNA degradation/decay and splicing pathways. (B) Differential gene expression analysis shows log2 of fold change of *MAPT mRNA* expression in the absence of *OTULIN* gene, and the STRING protein-protein interaction network analysis shows downregulation of predicted OTULIN-binding proteins encoding genes. Notably, the deubiquitinase OTULIN deficiency promotes the downregulation of genes associated with M1-linked linear ubiquitin ligase, LUBAC (*HOIP*, *HOIL-1,* and *SHARPIN*). (C) Differential gene expression analysis of ubiquitin pathway in OTULIN KO shows that the ubiquitin-modifying enzyme (E1), *UBA3* is upregulated along with a few ubiquitin-conjugating enzymes (E2s) and RNA-binding ubiquitin ligases (E3s), which are known to be involved in RNA catabolism. (D) Differential gene expression analysis of the RNA stability pathway shows that the CCR4-NOT (*CNOT*) complex is upregulated and can positively regulate mRNA degradation in *OTULIN* KO line. Many of the mRNA decay regulatory genes are also upregulated along with heterogeneous ribonuclear proteins (*HNRNPs*) and neurodegenerative disease-associated RNA-binding proteins.

## Discussion

We provide a previously unknown non-canonical function of OTULIN in regulating mRNA levels of tau and other proteins involved in proteostatic/RNA metabolic pathways. We initially hypothesized that because OTULIN is an M1-linked deubiquinase, its loss would lead to tau turnover via one of the two major proteostatic pathways: the proteasome and autophagy. However, our surprising result on the mRNA regulatory role of OTULIN suggests a likely transcriptional regulation and/or RNA metabolism role of OTULIN in neurons, which may or may not be dependent on its deubiquitinase activity.

Protein ubiquitination is a three-step post-translational modification that requires the sequential action of three enzymes: ubiquitin-activating enzyme (E1), ubiquitin-conjugating enzyme (E2), and ubiquitin ligase enzyme (E3)^47^. However, protein ubiquitination becomes complex when decoding the functions of differential ubiquitination on the same protein with various linkages, such as M1-, K6-, K11-, K27-, K29-, K33-, K48-, and K63-linked ubiquitin chains. It also adds further complexity to understanding the roles of homotypic, hetero-typic, linear, and branched ubiquitination with various linkages on the same protein^48,8^. Such a complex ubiquitination occurs on the pathological tau aggregates in the brains of patients with tauopathies. Notably, prior studies have suggested that PHFs from AD brains show M1-, K6-, K11-, K48-, and K63-linked ubiquitin chains via mass-spectrometry^8,13,14,15^. Tau has been shown as a substrate of 20S proteasome, 26S proteasome, chaperone-mediated autophagy, microautophagy, macroautophagy, and aggrephagy depending on the post-translational modifications, suggesting that ubiquitin can modify tau function and its half-life depending on the cellular contexts^8^. However, it is unclear why tau requires differential ubiquitination to undergo degradation. Most likely, the sequential failure of various proteostasis machineries adds differential linkages of ubiquitin chains, such as K63- and M1-linked polyubiquitin, which recruit ubiquitin-binding adaptors or receptors to activate transcription factors. These factors, in turn, trigger gene expression for cell survival, immune responses, and the replenishment of proteostasis machineries for efficient clearance of protein aggregates.

The linear ubiquitin assembly complex (LUBAC) is composed of RNF31/HOIP, RBCK1/HOIL-1, and SHARPIN, which function as an E3-ubiquitin ligase for M1-linked polyubiquitination on protein aggregates associated with various neurodegenerative diseases^10^. The deubiquitinase OTULIN specifically deubiquitinates M1-linked ubiquitin chains, such as those found in HOIL-1 and SHARPIN, whereas CYLD deubiquitinates both M1- and K63-linked ubiquitin chains but not the LUBAC^26,27,28,29^. OTULIN functions as a negative regulator of LUBAC, NF-κB, and autophagy^10,11,24,25^. We, therefore, aimed to understand the role of OTULIN in the clearance of tau aggregates in AD. We utilized sporadic AD2.1 iPSC line-derived neurons and the human neuroblastoma SH-SY5Y *OTULIN* KO line as model systems to investigate the role of OTULIN in tau clearance and differential gene expression. Our first RNA sequence analyses of WTC11 and sAD2.1 iPSNs indicated that over 4,000 genes and transcripts are differentially expressed in sporadic sAD2.1 line-derived neurons compared to healthy control WTC11 neurons. The most notable hits are the melanoma antigen genes *MAGEA and MAGEA2B*, which are known to positively regulate the ubiquitin ligase activity of RING domain proteins^49^. *MAGEA2B* and *MAGEA* are significantly downregulated in sAD2.1 neurons, suggesting a likely loss of a few unknown ubiquitin ligase activities, which may explain the elevated pathological tau levels in sAD2.1 neurons. However, the level of M1-linked LUBAC E3-ligase components, including *HOIP, HOIL-1, and SHARPIN, was* not significantly altered in sAD2.1 neurons. Since MAGEs interact with more than 50 RING-type E3 ubiquitin ligases^50^, a link between MAGEs and impaired tau clearance in AD needs further investigation.

Interestingly, significant downregulation of OTULIN long non-coding RNA (lncRNA) was detected in sAD2.1 neurons, which negatively correlated with elevated OTULIN protein levels. Since lncRNAs are intronic^51^, overlapping, or antisense to protein-coding mRNAs (exons) like microRNAs (miRNAs), the down-regulated OTULIN lncRNA may likely increase the level of OTULIN expression in sAD2.1 neurons. In a previous study, we observed an inverse correlation between elevated mRNA and reduced long non-coding RNA (lncRNA) or circular RNA (circRNA)^52^. Notably, increased OTULIN is positively correlating with elevated levels of total tau and phosphorylated tau at AT8 (p-S202/p-T205), AT180 (p-T231), and PHF-1 (p-S396/p-S404) epitopes, suggesting that OTULIN may regulate tau pathology.

To understand the role of the M1-linked deubiquitinase activity of OTULIN in tau pathology, we treated the sporadic sAD2.1 neurons with compound UC495. Inhibiting OTULIN deubiquitinase with UC495 decreases phosphorylated tau at the AT8 epitope but not at the AT180 and PHF-1 epitopes or total tau levels. Notably, inhibiting the deubiquitinase activity of OTULIN does not affect the level of OTULIN in sAD2.1 neurons, suggesting that inhibiting OTULIN deubiquitinase alone is sufficient to prevent tau pathology moderately. However, the exact mechanism by which phosphorylated tau at the AT8 epitope is decreased upon inhibiting the OTULIN deubiquitinase with UC495 is unknown. It is possible that the compound UC495 may promote partial clearance of pathological tau by restoring proteostasis activity, such as by enhancing the E3-ubiquitin ligase of LUBAC or stabilizing M1-linked ubiquitin chains on ATG13, a key regulator of autophagosome initiation^46^. Alternatively, OTULIN inhibition may also likely downregulate the activation of certain tau kinases, such as CDK5, GSK-3β, and MAPK, or any protein kinases that phosphorylate tau at S202 and T205 (AT8). In contrast to the effect of compound UC495 in sAD2.1 neurons, OTULIN inhibition with UC495 in SH-SY5Y cells significantly decreases OTULIN, ubiquitin, and total tau; consequently, phosphorylated tau was not detectable with AT8, AT180, and PHF-1 antibodies. Although the effect of compound UC495 is inconsistent with cell types such as sAD2.1 neurons and SH-SY5Y neuroblastoma, UC495 could be a potential therapeutic molecule that may alleviate tau pathology in tauopathy brains for future study.

To better understand the role of OTULIN in tau pathology, the *OTULIN* gene was knocked out in sAD2.1 neurons or the SH-SY5Y neuroblastoma line. We observed that the tau protein completely disappeared in both cell types of *OTULIN* KO. The overall morphology of sAD2.1 neurons was not affected in the absence of OTULIN; however, it does exhibit morphological defects, including a lack of neurite-like projections, in the majority of SH-SY5Y *OTULIN* KO cells. We assumed that the loss of tau in *OTULIN* KO is due to enhanced proteostasis activity. We, therefore, used various proteostasis inhibitors with protein translation inhibitor, cycloheximide (CHX), to understand the dynamics of tau degradation, including lactacystin (20S proteasome), MG132 (26S proteasome), Bafilomycin-A1 (Baf-A1; autophagy), and PD 150606 (Calpain 1/2). Surprisingly, none of the proteostasis inhibitors stabilize tau, which eventually motivated us to analyze MAPT mRNA levels by RT-qPCR, which suggests that *MAPT* mRNA has completely disappeared in the absence of OTULIN in the SH-SY5Y line. This finding further motivated us to perform bulk RNA sequencing to analyze whether the effects of *OTULIN* KO are specific to *MAPT* mRNA or dysregulated RNA metabolism globally.

Bulk RNA sequence analysis reveals that OTULIN KO dysregulates RNA metabolism by downregulating the expression of 13,341 genes and 43,003 transcripts and upregulating 774 genes and 1,113 transcripts, respectively, compared to SH-SY5Y WT. It confirms that the effect of *OTULIN* KO is not specific to the downregulation of the *MAPT* gene. Still, it does globally and most likely functions as a master regulator of RNA metabolism. However, the mechanism of OTULIN deficiency-associated dysregulated RNA metabolism is unknown. We, therefore, searched the possible candidates from our RNA sequencing data relevant to OTULIN-binding proteins, RNA metabolism associated with mRNA decay/degradation, mRNA stability, DNA transcription repressors, ubiquitination, and RNA-binding proteins. Primarily, we searched the differential expression of OTULIN-binding proteins obtained from the STRING protein-protein interactions database and identified that none of them were detectable at mRNA levels, including the ubiquitin pathway-associated LUBAC components (*HOIP*, *HOIL-1*, *SHARPIN*), *VCP*, Ubiquitin B (*UBB*) and Ubiquitin C (*UBC*), and Neural Precursor Cell Expressed, Developmentally Down-Regulated 8 (*NEDD8*).

Our RNA sequencing analysis showed that *OTULIN* KO in SH-SY5Y upregulated the expression of 774 genes and 1,113 transcripts that are most likely required for cell survival under autoimmunity by suppressing gene transcription and accelerating RNA catabolism. We identified upregulation of ubiquitin-like modifier enzyme (*UBA3*), which is a component of NEDD8-activating enzyme 1 (NAE1), and its associated neddylation has been shown as a regulator of neuronal aging and neurodegeneration in AD^53^. Many ubiquitin-conjugating enzymes (UBE2s), including *UBE2B/D2/D3/E3/H/K/L3/L5/M/N/Q1/V2/W* were upregulated among the approximately 40 E2 enzymes. However, many proteasome subunit genes were downregulated except proteasome activator subunit 3 (*PSME3*), proteasome 20S subunit beta 7 (*PSMB7*), proteasome 26S subunit, ATPase 6 (*PSMC6*), proteasome 26S subunit ubiquitin receptor, non-ATPase 2 (*PSMD2*), and proteasome 26S subunit, non-ATPase 14 (*PSMD14*) genes that were upregulated. Since more than 600 ubiquitin-ligases (E3s) are involved in the ubiquitin-proteasome pathway, we searched for RNA-binding ubiquitin ligases^54^. We identified that most of them are downregulated except Ring finger and CCCH-type domains 2 (*RC3H2*) and Mex-3 RNA binding family member C (*MEX3C*), which are known to bind to 3’-UTR of mRNAs, leading to deadenylation and degradation of mRNAs such as MHC-I mRNA^55^. Interestingly, we identified that many mRNA stability regulator proteins encoding genes such as Carbon catabolite repression 4-negative on TATA-less (CCR4-NOT) complex (*CNOT*)^56,57^, YTH N6-Methyladenosine RNA binding protein F (*YTHDF1/2/3*), Poly(A) specific ribonuclease subunit 2 (*PAN2*), and *PAN3* were either up or downregulated in *OTULIN* KO cells that might have regulated mRNA stability or degradation depending on the context. In addition to accelerating mRNA degradation/decay, *OTULIN* KO may suppress DNA transcription globally by upregulating the expression of transcription factors such as Yin Yang 1 (YY1) and Specificity protein 3 (SP3), which are known to function as suppressors of gene expression. *OTULIN* KO also upregulates many heterogeneous ribonucleoproteins (*HNRNPs*) and neurodegenerative disease-associated RNA-binding proteins such as Fragile X messenger ribonucleoprotein 1 (FMR1), TAR DNA binding protein (TARDBP/TDP-43), Ataxin 2 (ATXN2), and Musashi RNA binding protein 1 (MSI1).

In conclusion, the canonical function of OTULIN is counteracting the LUBAC (RNF31/HOIP, RBCK1/HOIL-1, and SHARPIN) E3-ubiquitin ligase leading to destabilization of LUBAC by deubiquitinating the linear M1-linked polyubiquitination on HOIL-1 and SHARPIN. The deubiquitinase of OTULIN functions as a negative regulator of LUBAC and M1-linked ubiquitin chains-associated NF-κB and autophagy. Knocking out of *OTULIN* in SH-SY5Y hyperactivates LUBAC, leading to increased M1-linked linear polyubiquitination and autoimmunity, which in turn negatively regulates LUBAC activity by downregulating its components and many thousands of genes globally to prevent autoimmunity for cell survival. We recognized the non-canonical function of OTULIN as a master regulator of tau encoding *MAPT* mRNA expression and RNA metabolism, significantly impacting dysregulated RNA metabolism in various neurodegenerative diseases.

## Materials and Methods

### Cell culture

The WTC11 iPSCs^58^ were a kind gift from Dr. Li Gan and sAD2.1 iPSCs^48^ were obtained from Coriell^®^. Neural progenitor cells (NPCs) of WTC11 and sAD2.1 were first cultured in STEMdiff forebrain neuron differentiation kit (STEMCELL Tech # 08600) and then matured in BrainPhys neuronal medium with SM1 supplement (STEMCELL Tech # 05792). HEK293T cells were cultured in DMEM (ThermoFisher # 11965092) and supplemented with 10% heat-inactivated fetal bovine serum (FBS) (ThermoFisher # 16140071) and 2 mM L-Glutamine (ThermoFisher # 25030081). All the cell cultures were maintained at 37° C with 5% CO2 and moisture. SH-SY5Y cells were grown in DMEM-F12 (1:1) (ThermoFisher # 11320033) culture medium supplemented with 10% FBS and 1X concentration of penicillin-streptomycin (10,000 U/mL, stock) (ThermoFisher # 15140122) for UC495 treatment and *OTULIN* knockout.

### Lentivirus production

The day prior to lentiviral transduction of single guide (sg) RNA of *OTULIN*, HEK293T cells were plated onto a 6-well plate at a 70% confluency. 60µl (6µg) of the single guide RNA to OTULIN (sgOTULIN) plasmid^32^ was added to 30µl (3µg) of psPAX packaging plasmid DNA and 15µl (1.5µg) of pVSV-G envelope plasmid DNA in 345µl of nuclease-free water with 50µl of 2M CaCl2. The DNA mix was then added dropwise to 500µl of 2x HBS buffer while slowly vertexing and incubated at room temperature for 30 minutes. Then, the DNA mix solution was added to the HEK293T and incubated overnight at 37° C. The culture of media was changed 24 hours later. The conditioned media was harvested 48 hours later and passed through a 0.2µm syringe filter. The lentivirus was then aliquoted and kept frozen at −80° C.

### Lentivirus transduction of neural progenitor cells (NPCs)

Thawed Lentivirus (100µl) was added to WTC11, and sAD2.1 NPCs were cultured in STEMdiff media onto a 6-well plate pre-coated with 200µl Geltrex (ThermoFisher # A1569601). After overnight incubation, the lentivirus was aspirated and replaced with fresh media. 0.5 µg/ml puromycin (ThermoFisher # J67236.XF) was added 48hrs later for selection. After the cells reached a 70% confluency in media + puromycin, they were moved to a 6-well plate pre-coated with 10µg Poly-L-Ornithine (Sigma # P4957-50ML) and 15µg Laminin (Sigma # L6274-.5MG) and fed with BrainPhys neuronal media (STEMCELL # 05790) for 20 days. Cells were then harvested and lysed in cold RIPA buffer (ThermoFisher # 89901) with 1% PMSF (Sigma # 93482-50ML-F), 1X concentration of protease (Sigma # P8340-1ML) and phosphatase (Sigma # P2850-1ML) inhibitor cocktails. Lysates were spun down at 14,000g for 20 minutes at 4° C. Lysates were transferred to new collection tubes and stored at −80° C for further analysis.

### Inhibition of deubiquitinase OTULIN with compound UC495

Compound UC495 was originally identified through a virtual screening effort to discover small molecule OTULIN inhibitors and subsequently synthesized. 1µM of UC495 in DMSO was added to matured sporadic sAD2.1 iPSC line-derived neurons and incubated overnight in a cell culture incubator. 2µM of UC495 in DMSO was added to the SH-SY5Y cells and incubated overnight. After the UC495 treatment, cells were harvested for western blotting.

### Bulk RNA sequencing

RNA was extracted from healthy control WTC11 iPSC line- and sporadic sAD2.1 iPSC line-derived neurons, and human neuroblastoma SH-SY5Y WT as control and SH-SY5Y OUTLIN KO, respectively. RNA was extracted with Trizol reagent (ThermoFisher # 15596026) and quantified by NanoDrop. 2.0 mg of RNA was sent to LC Sciences, and LC Sciences performed poly(A) RNA-Sequencing - sample quality control, library preparation, sequencing (150 bp PE, 6GB data per sample), and analysis. *De Novo* assembly, alignment of RNA-Seq reads to reference genome, identification and construction of splice-junctions, report of known and novel transcripts with annotation and abundance, identification of alternate splicing and report of isoform abundance, test for differential expression at gene level and transcript level, Gene Ontology (GO) and Kyoto Encyclopedia of Genes and Genomes (KEGG) annotation and enrichment analysis Gene Set Enrichment Analyses (GSEA) were done by the LC Sciences. All raw RNA sequence analysis data was deposited to NCBI GEO with accession # xxxxx (WTC11/sAD2.1 iPSNs) and # xxxxx (SH-SY5Y WT/OTULIN KO).

### Proteostasis inhibitors treatment and western blotting

Human neuroblastoma SH-SY5Y WT/*OTULIN* KO cells (0.1 × 10^6^) were seeded in a 12-well plate. After overnight culturing of cells, protein translation was inhibited with 50µM of cycloheximide (CHX) 1h prior to adding the proteostasis inhibitors, 10µM MG132 (26S proteasome), 10µM lactacystin (20S proteasome), 100 nM Bafilomycin A1 (Baf A1, autophagy), 100µM PD 150606 (calpain 1/2) either individually or all four inhibitors together for 4h. All the inhibitors were reconstituted with DMSO to prepare stock solutions and stored at −20° C until further use. After treatment, cells were washed with 1X PBS three times and lysed with ice-cold RIPA buffer with PMSF, protease, and phosphatase inhibitor cocktails and incubated on a rocker for 30min at 4° C. Then the cell lysates were centrifuged at 14,000g for 10min at 4° C. The resulting supernatant was stored at −80° C or heat-denatured the lysates with 1X LDS-sample buffer (ThermoFisher # NP0007) at 95° C for 10min. Prior to loading the denatured samples onto a gel for protein separation, samples were centrifuged to collect the supernatant at 14,000g for 5min at 4° C.

### Proteostasis inhibitors and antibodies

The following proteostasis inhibitors were procured from Cayman Chemical. Cycloheximide (Cayman # 26924), Lactacystin (Cayman # 70980), MG132 (Cayman # 13697), Baf A1 (Cayman # 11038), and PD 150606 (Cayman # 13859). The following antibodies were procured from various vendors and their dilution mentioned here. β-Actin, Rb mAb (1:1000; CST # 4970S), OLTULIN, Rb pAb (1:1000; CST # 14127S) Ubiquitin, Rb pAb (CST # 58395S), Tau12, Mo mAb (1:2000; Sigma # MAB2241), AT8, Mo mAb (1: 8000; ThermoFisher # MN1020), AT180, Mo mAb (1:8000, ThermoFisher # MN1040), PHF-1 (1:10,000, a kind gift from Dr. Peter Davies^59^), and NeuN (1:1500; GeneTex # GTX638922).

## Online supplemental materials

### Author Contributions

K.B., and F.F.L – developed the project and acquired funding. K.T – conceived OTULIN is a master regulator of gene expression and RNA metabolism. K.T., F.F.L., and K.B – designed the experiments. K.T., V.B – performed the experiments and analyzed the data under the supervision of K.B. V.B – responsible for sAD2.1 OTULIN KO. K.T – responsible for SH-SY5Y OTULIN KO. M.L., and W.L – responsible for the identification and synthesis of UC495 compound. K.T., and K.B – wrote the manuscript with input from all authors. Authors have read and agreed to publish the manuscript.

### Funding

This research work was funded by the National Institutes of Health (NIH) funding: (1) RF1AG072703 (to F.F.L). (2) RF1NS083704-05A1, R01NS083704, New Mexico Higher Education Department – Technology Enhancement Fund (TEF), RF1AG072703-01A1, University of New Mexico (UNM) Health Sciences Center Bridge Funding, UNM Department of Molecular Genetics and Microbiology intradepartmental grant, the New Mexico Alzheimer’s Disease Research Center (NM ADRC) P30 grant P30AG086404-01 funding (to K.B.); (3) UNM Center for Biomedical Research Excellence (CoBRE) in Center for Brain Recovery and Repair Pre-Clinical Core P20GM109089. Autophagy, Inflammation, and Metabolism (AIM) CoBRE Center P20GM121176-04. (4) A pilot development grant award from P30 parent grant P30AG086404-01 (to K.T).

## Acknowledgments

We thank Dr. Li Gan, Weill Cornell Medical Center for providing the WTC11 iPSC line.

