## Supplemental Information for "The deubiquitinase OTULIN regulates tau expression and RNA metabolism in neurons"

**Table S1: Reads quality control (WTC11 vs sAD2.1)**

| Sample | Raw Data | | Valid Data | | Valid Ratio(reads) | Q20% | Q30% | GC content% |
|---|---|---|---|---|---|---|---|---|
| | Read | Base | Read | Base | | | | |
| sAD_1 | 41327090 | 6.20G | 40051318 | 6.01G | 96.91 | 99.97 | 96.92 | 47.50 |
| sAD_2 | 44583144 | 6.69G | 43377306 | 6.51G | 97.30 | 99.97 | 96.56 | 47.50 |
| sAD_3 | 40911758 | 6.14G | 39517352 | 5.93G | 96.59 | 99.96 | 96.71 | 48.50 |
| WTCII_1 | 40062948 | 6.01G | 38553188 | 5.78G | 96.23 | 99.96 | 96.40 | 46.50 |
| WTCII_2 | 50406868 | 7.56G | 46293998 | 6.94G | 91.84 | 99.95 | 96.15 | 49.50 |
| WTCII_3 | 39730098 | 5.96G | 38223934 | 5.73G | 96.21 | 99.96 | 96.48 | 46.50 |

**Table S2: 3D mapped region statistics (WTC11 vs sAD2.1)**

| Samples | sAD_1 | sAD_2 | sAD_3 | WTCII_1 | WTCII_2 | WTCII_3 |
|---|---|---|---|---|---|---|
| exon | 95.46 | 93.64 | 93.68 | 88.10 | 91.87 | 89.01 |
| intron | 4.11 | 5.89 | 5.85 | 10.95 | 7.39 | 10.12 |
| intergenic | 0.43 | 0.47 | 0.47 | 0.95 | 0.74 | 0.87 |

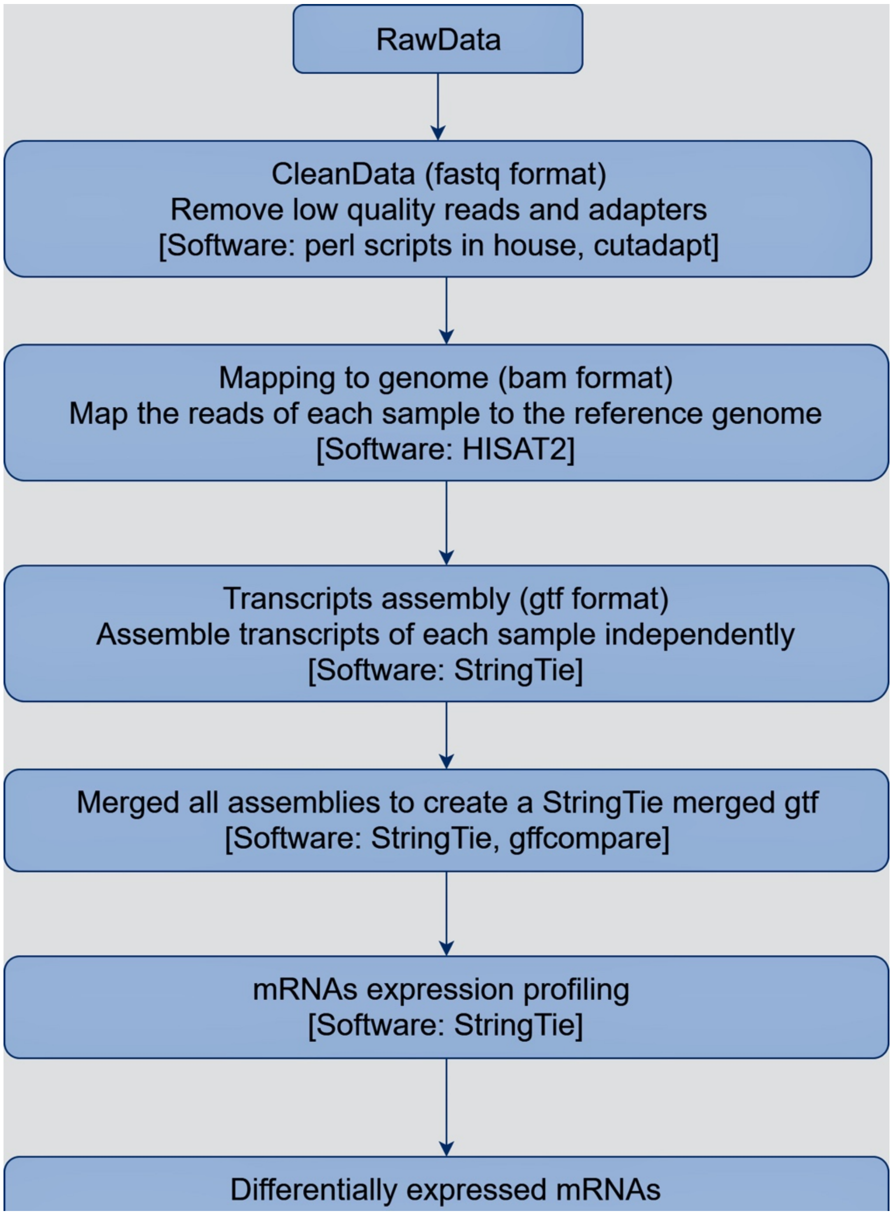


**Figure S1: Pipeline for the RNA sequence analyses in WTC11 and sAD2.1 iPSC-derived neurons (iPSNs).**

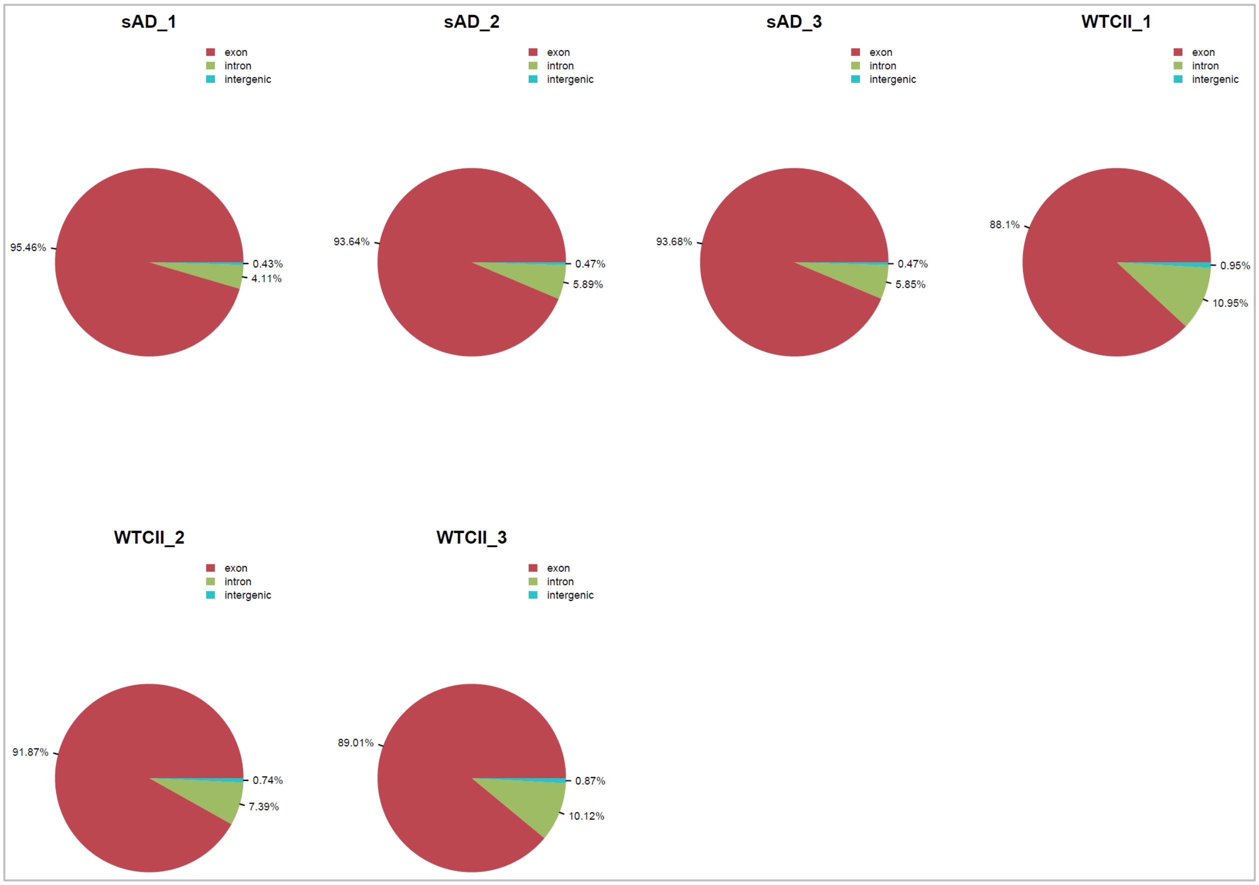


**Figure S2: Statistics of the mapped regions across six different samples.**

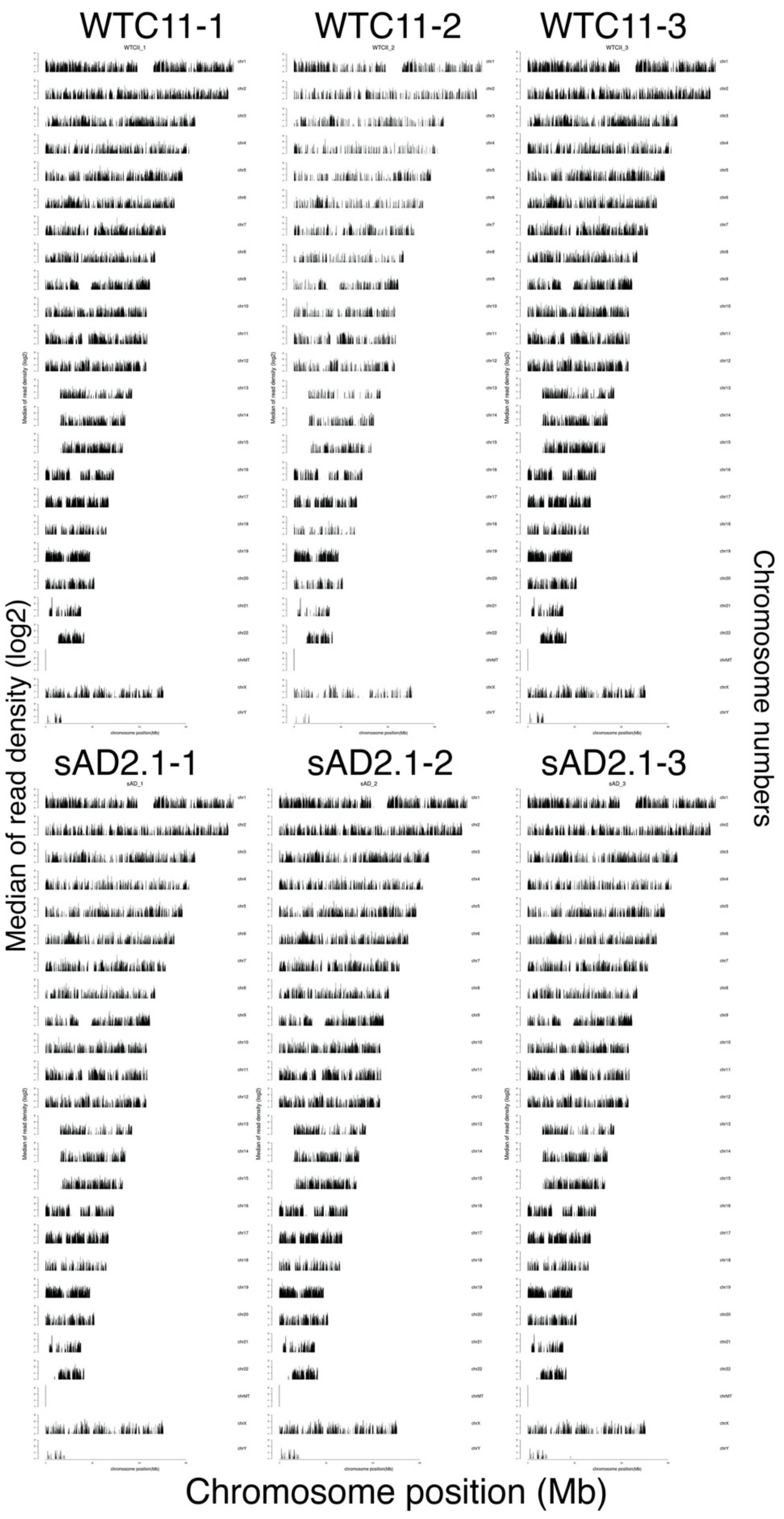


**Figure S3: Chromosome position and median read density distribution for six different samples.**

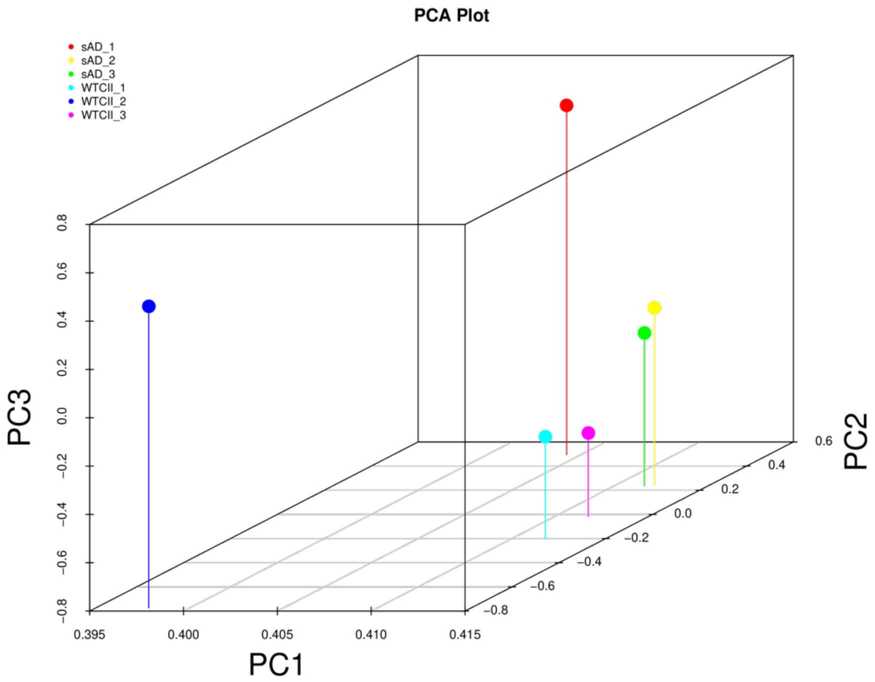


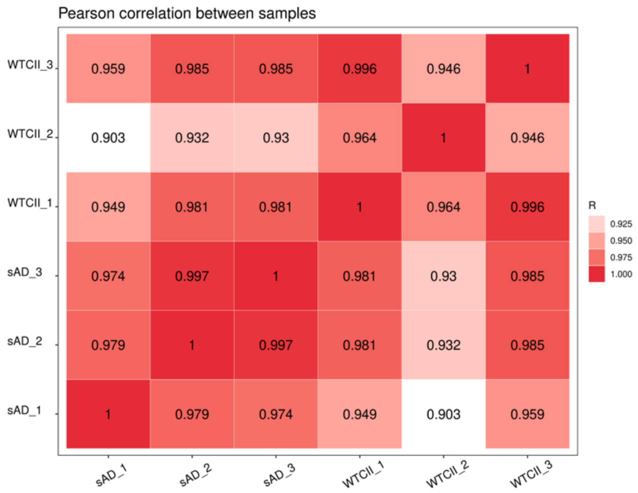


**Figure S4: Principal component analyses and correlation matrix for all six samples.**

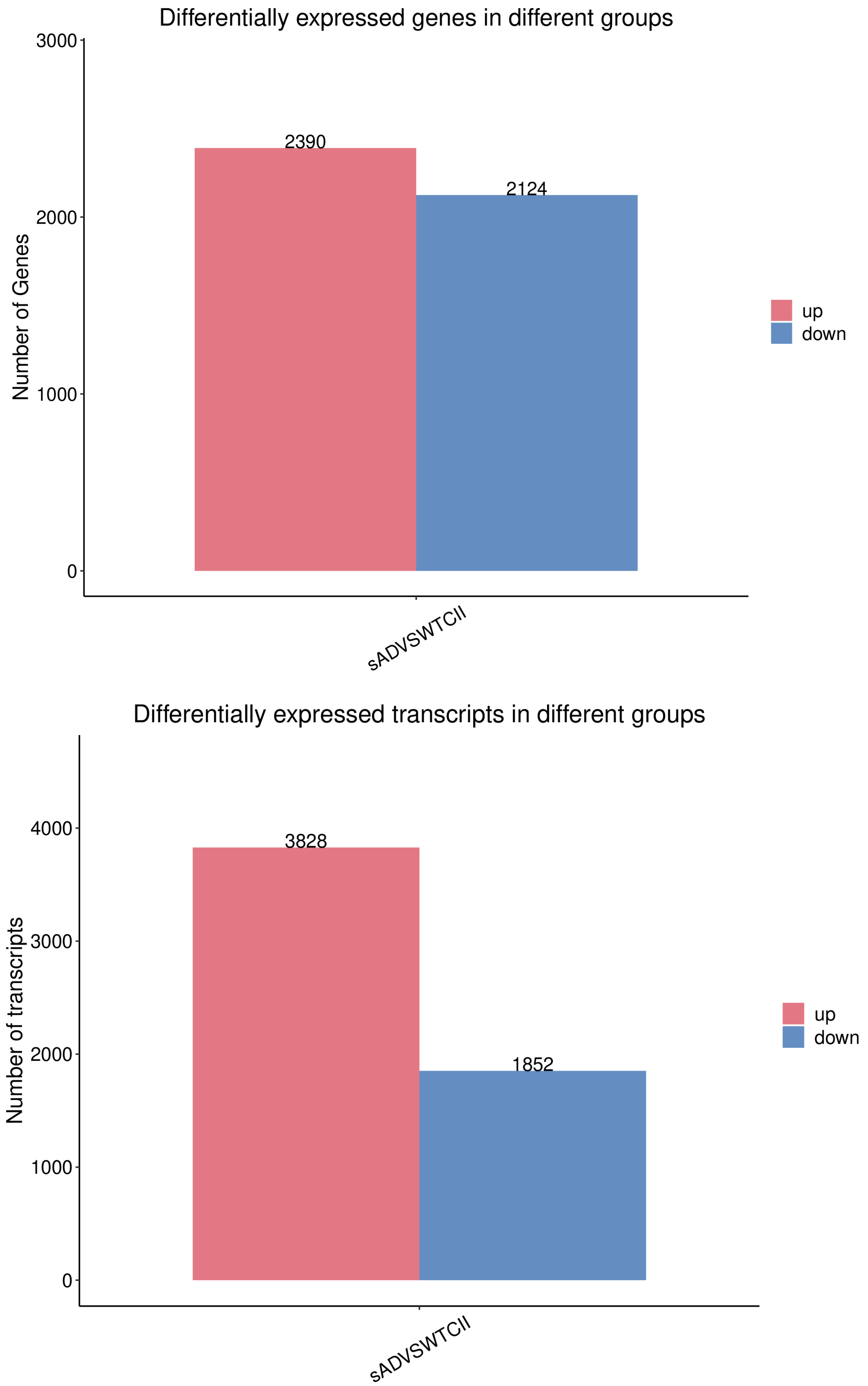


**Figure S5: Number of differentially expressed genes and transcripts in WTC11 vs sAD2.1 iPSNs.**

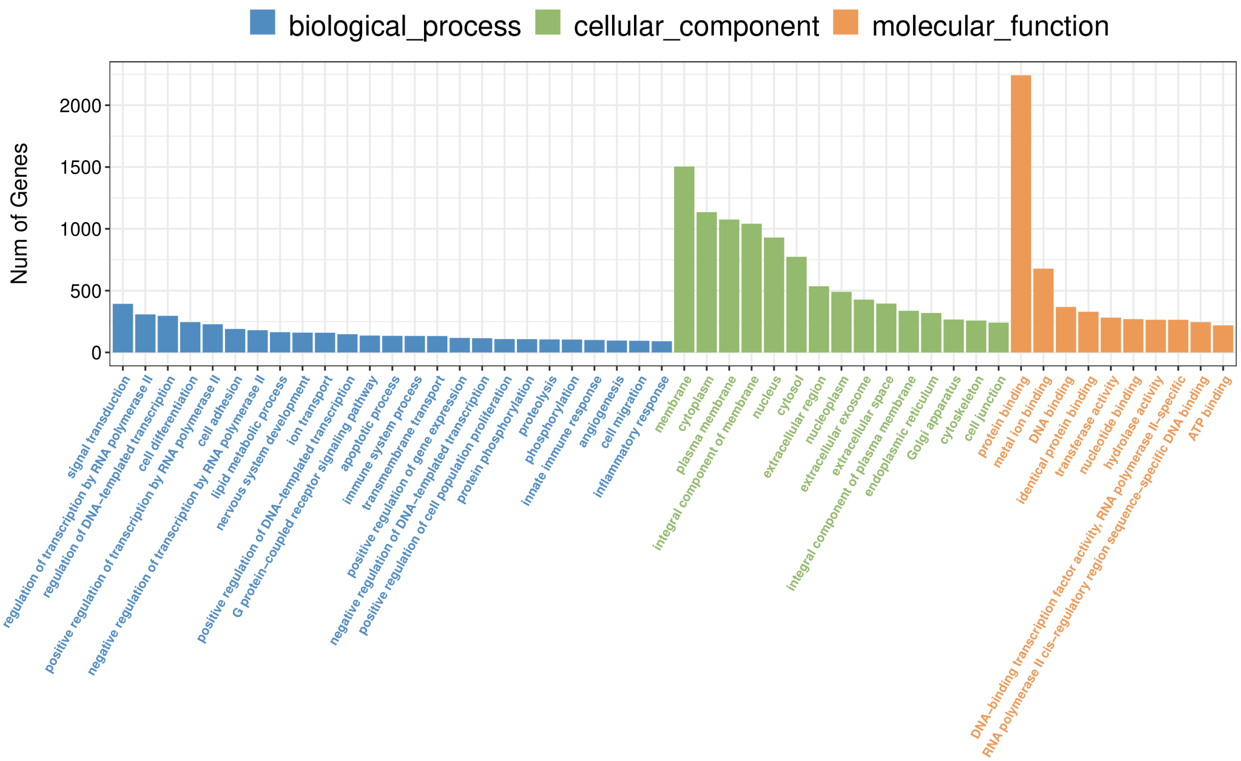


**Figure S6: Gene ontology analyses of differentially expressed genes in sAD2.1 iPSNs.** Gene ontology analyses of the association of number of differentially expressed genes in WTC11 vs sAD2.1 iPSNs relevant to biological processes, cellular component and molecular function. Note that protein binding represented highest number of differentially altered genes in sAD2.1 compared to WTC11.

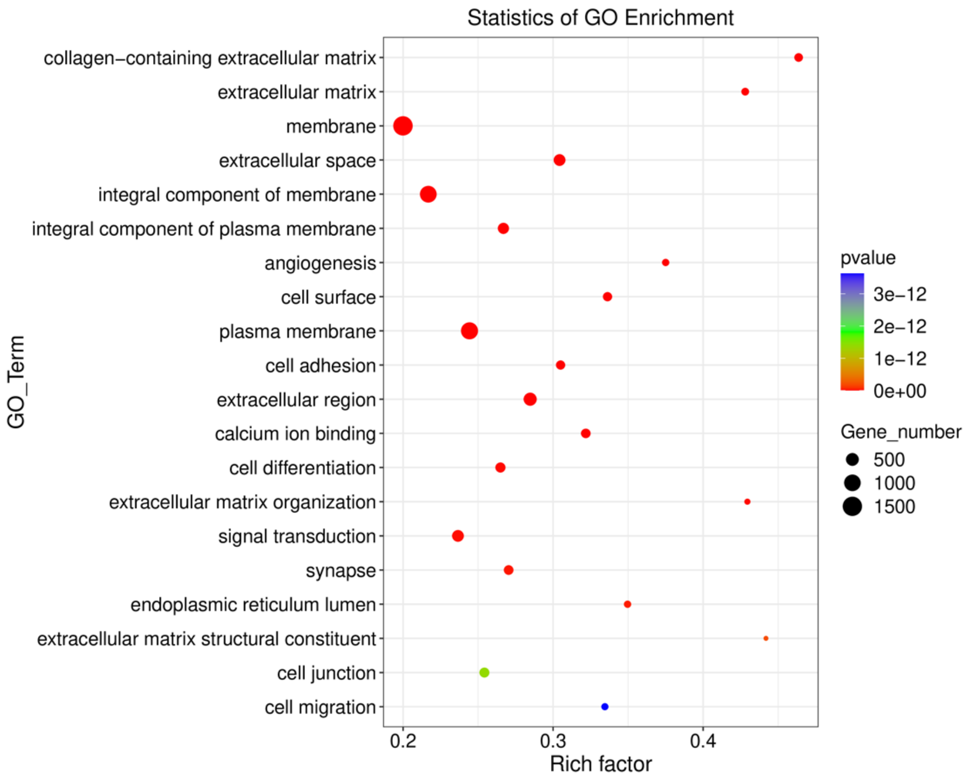


**Figure S7: Gene ontology enrichment plots in sAD2.1 vs WTC11 iPSNs.** Gene ontology enrichment plots showing number of genes associated with major cellular pathways.

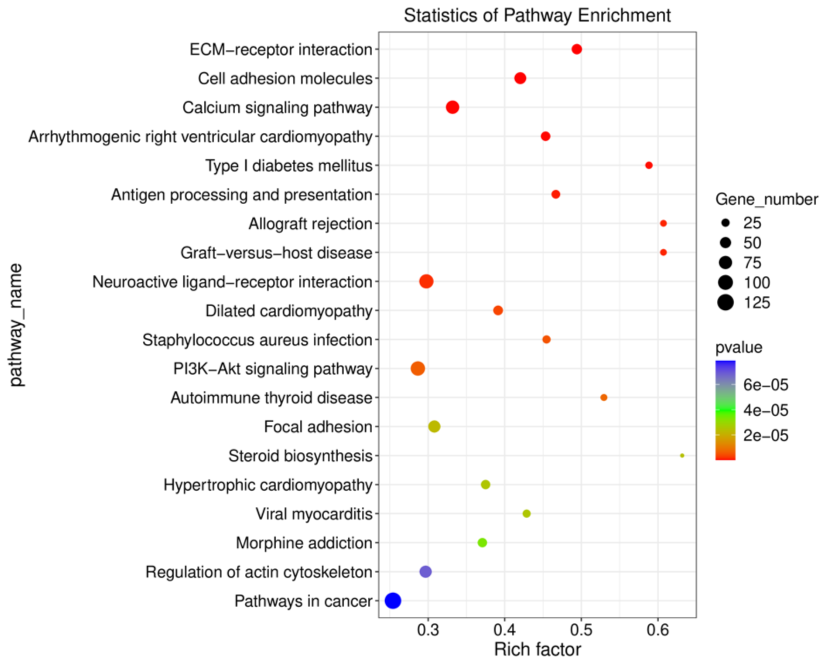


**Figure S8: KEGG enrichment plots in sAD2.1 vs WTC11 iPSNs.** KEGG enrichment plots show several genes associated with major cellular pathways.

**Table S3: Reads quality control (SH-SY5Y KO vs WT)**

| Sample | Raw Data | | Valid Data | | Valid Ratio(reads) | Q20% | Q30% | GC content% |
|---|---|---|---|---|---|---|---|---|
| | Read | Base | Read | Base | | | | |
| KO_1 | 38817316 | 5.82G | 37787998 | 5.67G | 97.35 | 99.75 | 97.74 | 48.50 |
| KO_2 | 37457530 | 5.62G | 36486542 | 5.47G | 97.41 | 99.75 | 97.46 | 48.50 |
| KO_3 | 38730082 | 5.81G | 37730190 | 5.66G | 97.42 | 99.77 | 97.87 | 48.50 |
| WT_1 | 41117518 | 6.17G | 39704186 | 5.96G | 96.56 | 99.75 | 97.61 | 49.50 |
| WT_2 | 41528248 | 7.56G | 40190316 | 6.03G | 96.78 | 99.76 | 97.72 | 48.50 |
| WT_3 | 40040468 | 6.01G | 38765676 | 5.81G | 96.82 | 99.76 | 97.68 | 48.50 |

**Table S4: 3D mapped region statistics (SH-SY5Y OTULIN KO vs WT)**

| Samples | KO_1 | KO_2 | KO_3 | WT_1 | WT_2 | WT_3 |
|---|---|---|---|---|---|---|
| exon | 70.19 | 59.67 | 78.12 | 92.75 | 94.16 | 93.98 |
| intron | 29.10 | 39.65 | 21.26 | 6.82 | 5.43 | 5.61 |
| intergenic | 0.71 | 0.68 | 0.61 | 0.42 | 0.41 | 0.41 |

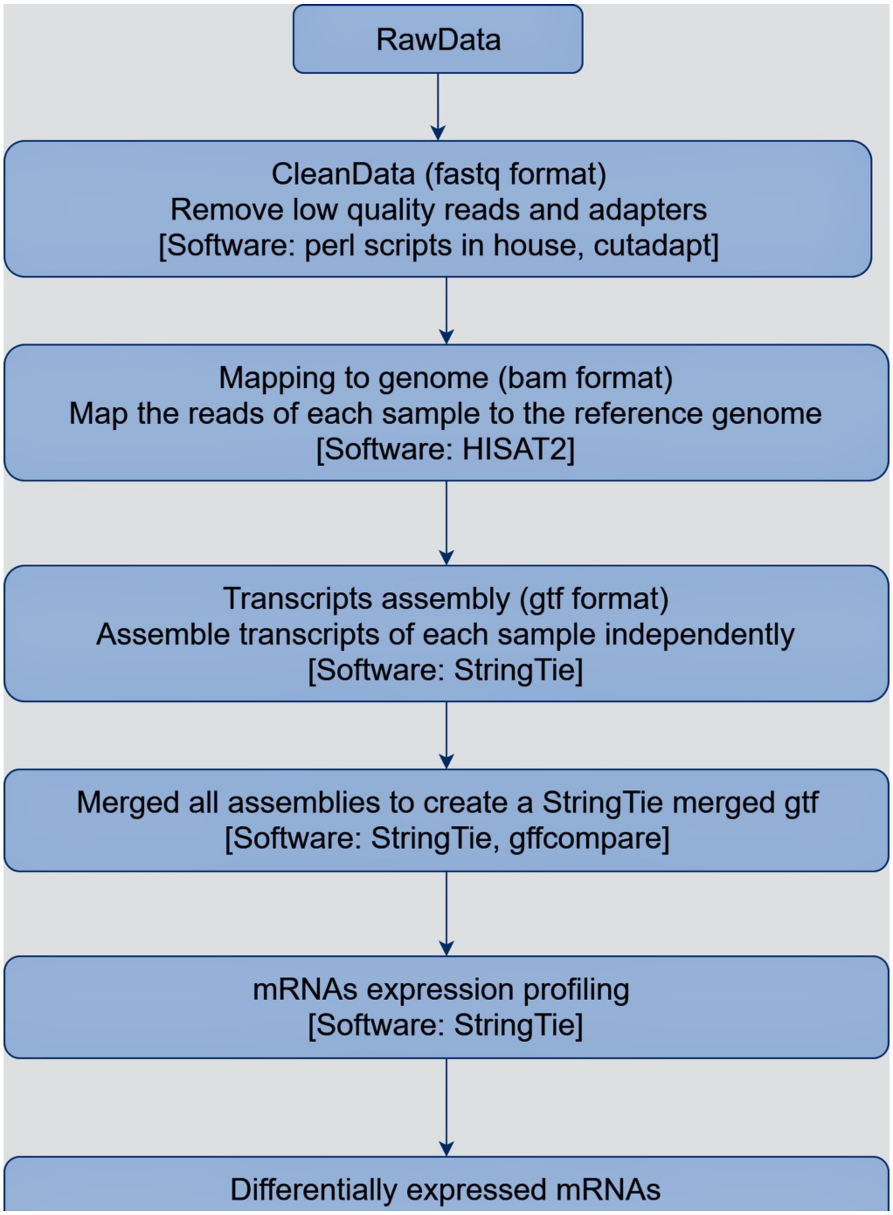


**Figure S9: Pipeline for the RNA sequence analyses in SH-SY5Y *OTULIN* KO and wild type (WT)**

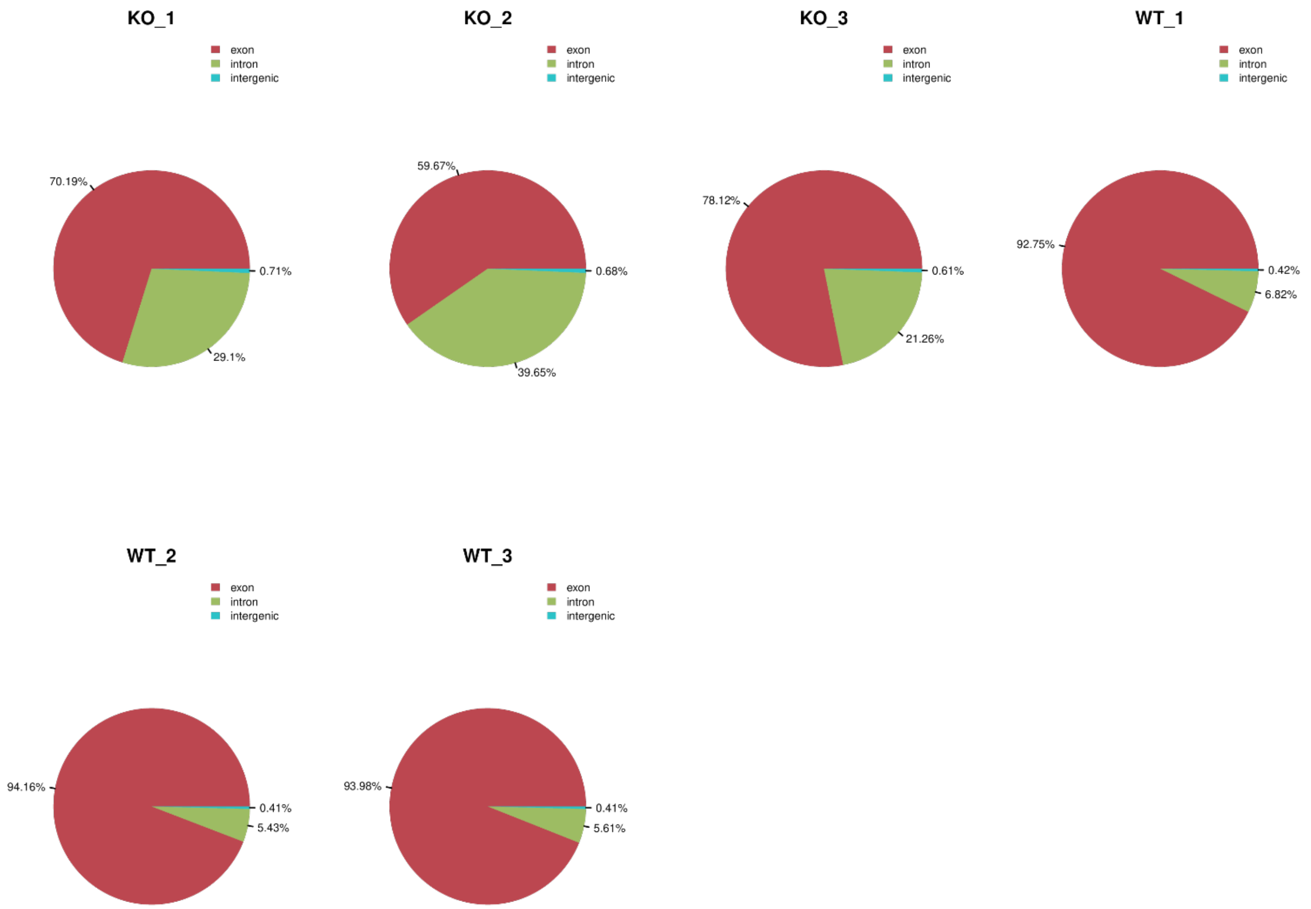


**Figure S10: Statistics of the mapped regions across six different samples (SH-SY5Y OTULIN KO vs WT).**

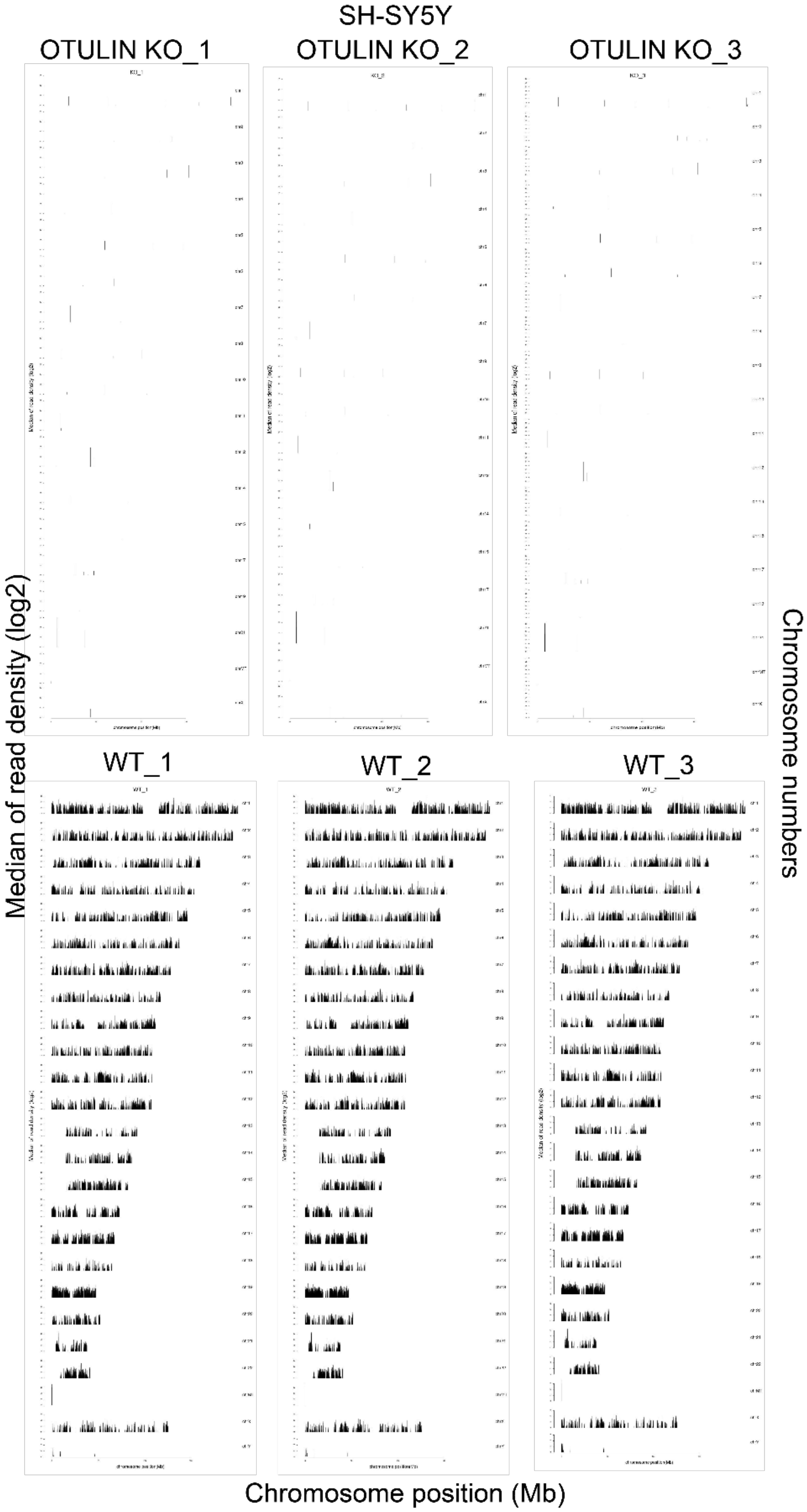


**Figure S11: Chromosome position and median read density distribution for six different samples (SH-SY5Y OTULIN KO vs WT).**

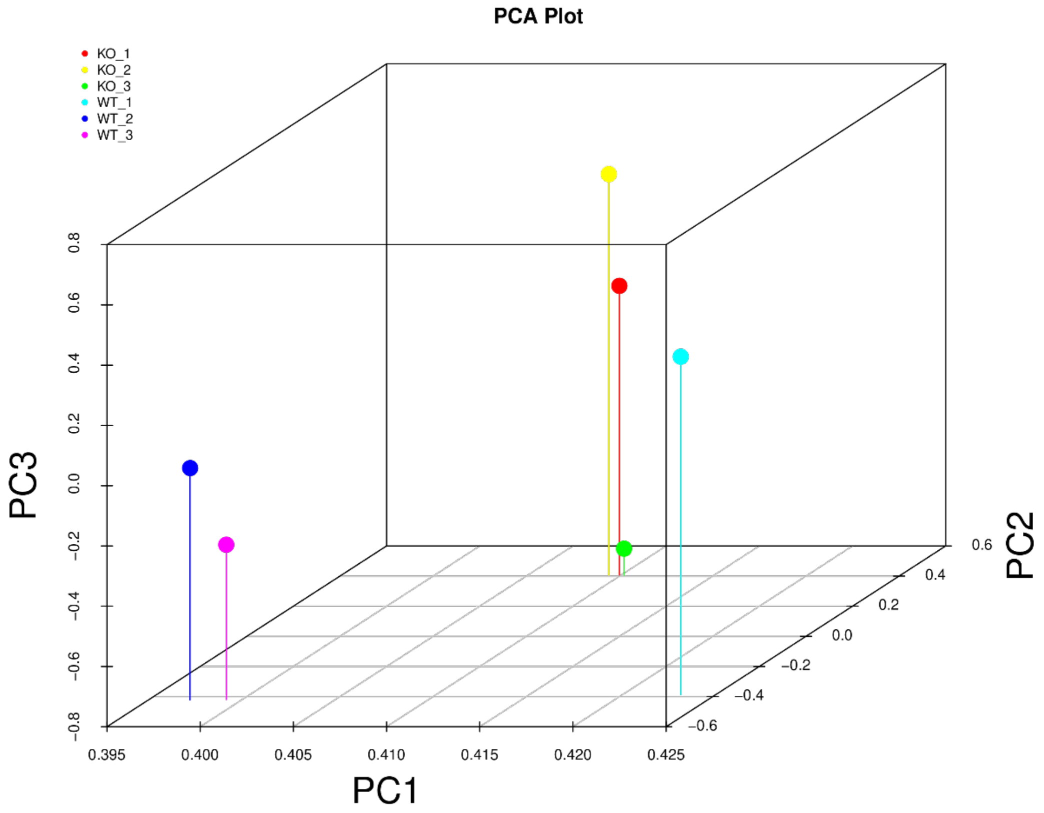


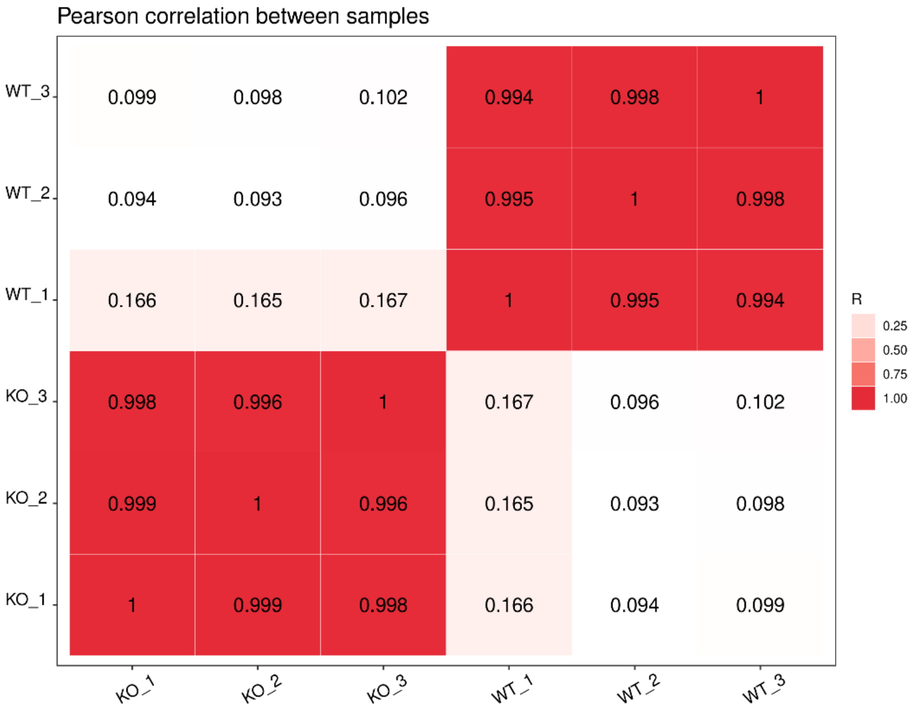


**Figure S12: Principal component analyses and correlation matrix for all six samples (SH-SY5Y OTULIN KO vs WT).**

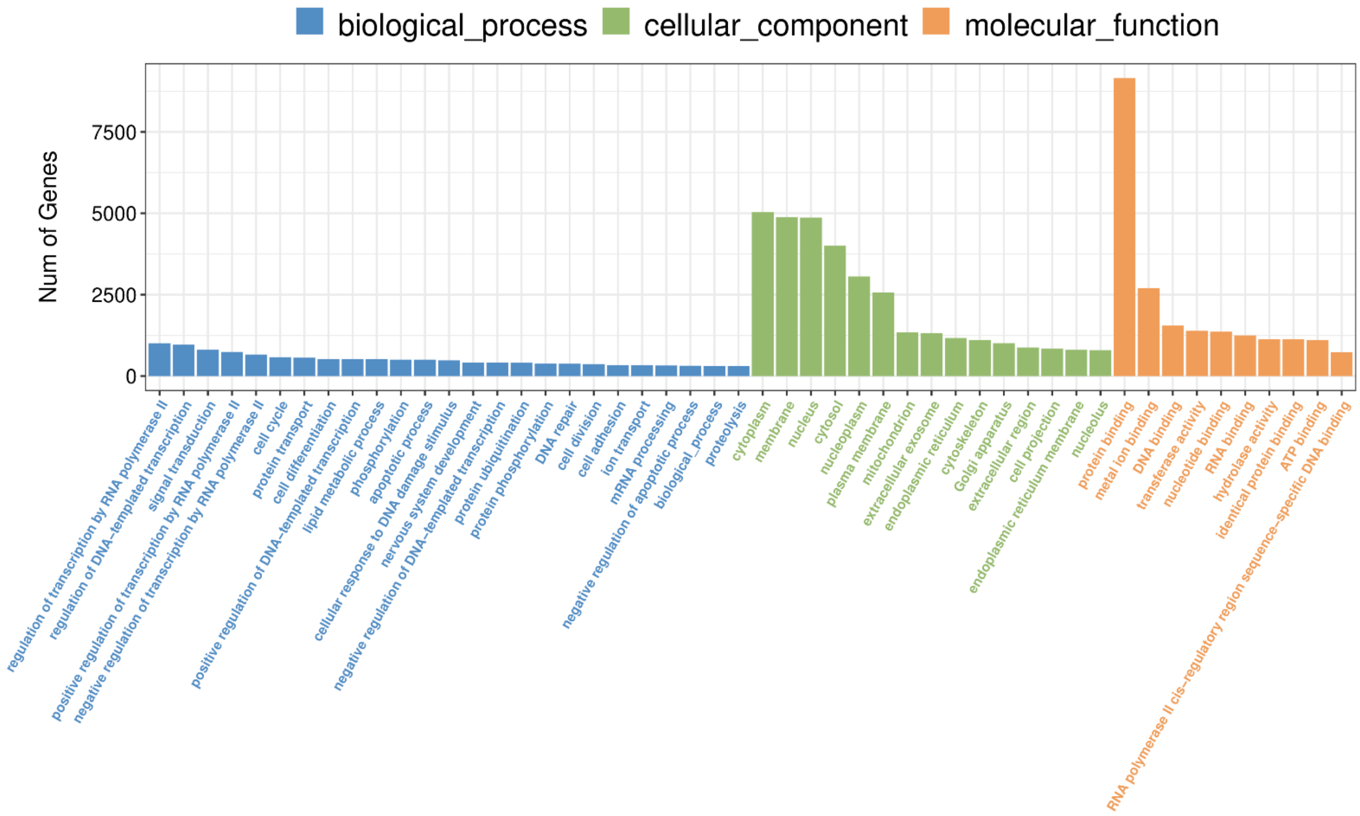


**Figure S13: Gene ontology analyses of differentially expressed genes in SH-SY5Y *OTULIN* KO.** Gene ontology analyses of the association of number of differentially expressed genes in *OTULIN* KO vs wild type SH-SY5Y relevant to biological processes, cellular component and molecular function. Note that protein binding represented highest number of differentially altered genes in OTULIN KO compared to wild type SH-SY5Y.

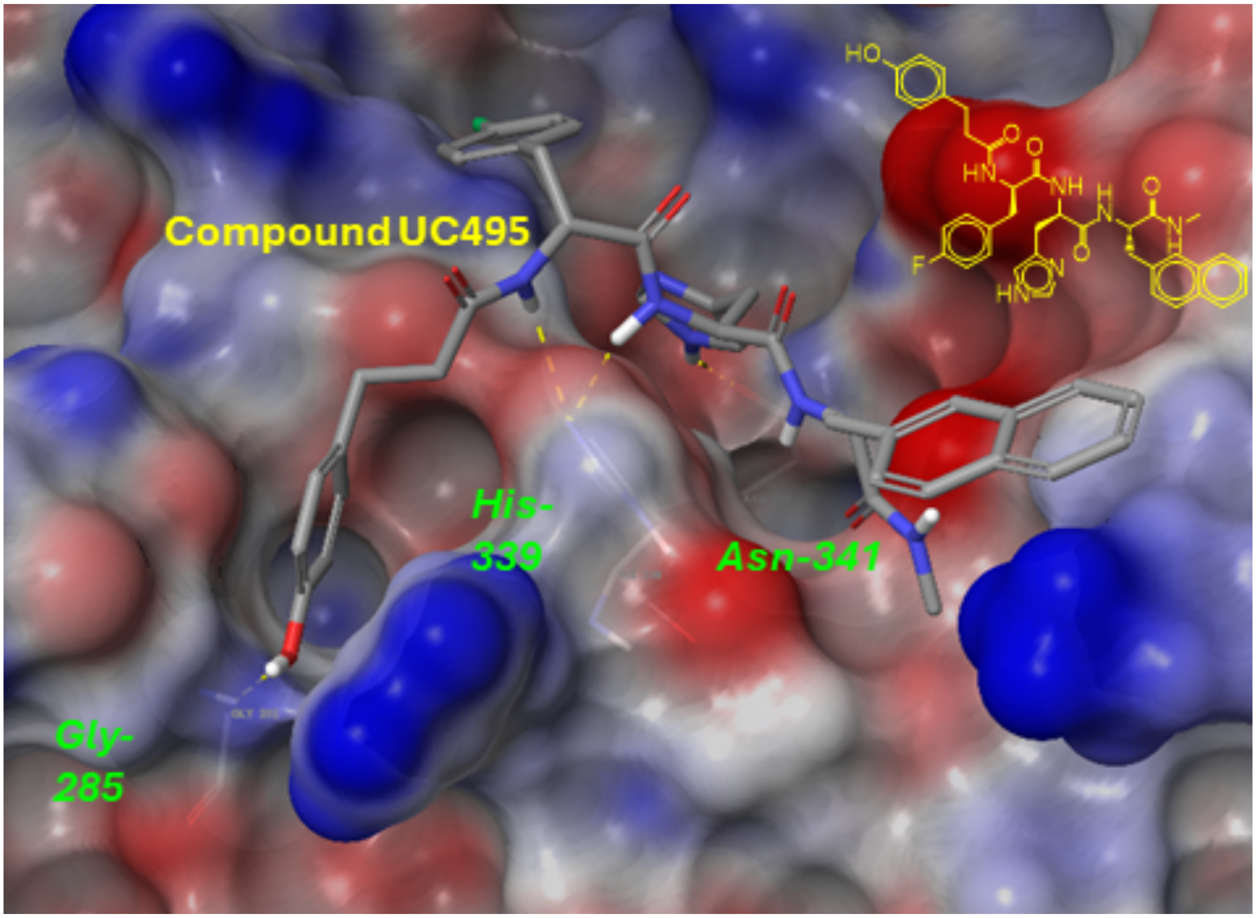


**Figure S14: In silico docking of UC495 with the deubiquitinase OTULIN catalytic center.** The identification of UC495 as an OTULIN inhibitor through a virtual screen. Compounds in a drug-like library were docked to the OTULIN catalytic center (PDB: 3ZNV) using the Schrodinger Molecule Modeling Suite. Major interactions of UC495 with OTULIN are shown in the figure. The OUTLIN surface is shown with electrostatic potentials, with red color indicating regions of high negative potential (electron-rich), blue color indicating regions of high positive potential (electron-poor), and other colors representing intermediate potentials.
